# Preserving functional network structure under anesthesia in the marmoset monkey brain

**DOI:** 10.1101/2023.11.21.568138

**Authors:** Michael Ortiz-Rios, Nikoloz Sirmpilatze, Jessica König, Susann Boreitus

## Abstract

Initiatives towards acquiring large-scale neuroimaging data in non-human primates promise improving translational neuroscience and cross-species comparisons. Crucial among these efforts is the need to expand sample sizes while reducing the impact of anesthesia on the functional properties of brain networks. Yet, the effects of anesthesia on non-human primate brain networks remains unclear. Here, we demonstrate that isoflurane anesthesia induces a variety of brain states in the marmoset brain with dramatically altered functional connectivity profiles. As an alternative, we recommend using a continuous infusion of the sedative medetomidine, supplemented with a low concentration of isoflurane. With this protocol, we observed robust visual activation during flickering light stimulation and identified resting-state networks similar to the awake state. In contrast, isoflurane alone led to a suppressed visual activation and the absence of awake-like network patterns. Comparing states using a graph-theoretical approach, we confirmed that the structure of functional networks is preserved under our proposed anesthesia protocol but is lost using isoflurane alone. We believe that the widespread adoption of this protocol will be a step towards advancing translational neuroscience initiatives in non-human primate neuroimaging. To promote the shared use of neuroimaging resources, we share our datasets on the Marmoset connectome project.

**Highlights:**

- Medetomidine combined with isoflurane is an effective protocol for driving visual BOLD responses compared to isoflurane-only anesthesia.
- Independent component analysis revealed similar resting-state networks for medetomidine-isoflurane anesthesia and awake states.
- The structure of resting-state networks was maintained under medetomidine-isoflurane but lost under isoflurane-only anesthesia.

## Introduction

Initiatives to gather extensive neuroimaging data in non-human primates (NHPs) aim to advance translational neuroscience and enable more accurate cross-species comparisons (Milham et al., 2020, 2018). Yet, translating findings across species faces significant methodological obstacles. A primary concern is whether NHP neuroimaging studies are conducted under awake or anesthetized states. Awake conditions are preferable for comparisons with humans, but training NHPs for awake neuroimaging remains logistically challenging (Schaeffer et al., 2022). Consequently, awake neuroimaging in NHPs is typically limited to a few animals (Milham et al., 2018; Ortiz-Rios et al., 2021). Conversely, using anesthesia is more practically feasible, allowing for larger study samples. However, anesthesia can disrupt brain network connectivity (Hutchison et al., 2014), complicating cross-species analysis. Thus, an anesthesia protocol that could preserve functional connectivity similar to the awake state can enhance the translational impact of cross-species comparisons by minimizing the negative effects of anesthesia and increasing the feasibility of large-scale neuroimaging studies in NHPs.

In NHPs, most neuroimaging data is acquired under anesthesia, commonly induced and maintained with isoflurane at concentration levels of 1% or higher (Milham et al., 2020, 2018). Isoflurane, however, negatively affects functional connectivity and reduces the probability of detecting resting state networks (RSNs) (Hori et al., 2020; Hutchison et al., 2014). Moreover, under isoflurane, the brain of both humans and NHPs may enter the burst-suppression state, a state that consists of quasi-periodic alternations between bursts of activity and periods of relative silence (Golkowski et al., 2017; Sirmpilatze et al., 2022; Zhang et al., 2019). We previously showed that burst-suppression results in a profound blood oxygen level dependent (BOLD) signal synchronization across the striatum and extensive regions of the neocortex in primates and rodents (Sirmpilatze et al., 2022). Additionally, lower concentrations of isoflurane typically induce continuous slow-wave activity, whereas concentrations exceeding those related to burst-suppression result in persistent suppression of cortical activity (Kroeger et al., 2013; Kroeger and Amzica, 2007). Therefore, datasets acquired during isoflurane anesthesia might contain a variety of brain states, each potentially exerting different effects on RSN patterns. This variability may produce contradictory results within and between studies and complicate comparisons with awake human fMRI studies.

To overcome the confounding effects of isoflurane anesthesia on fMRI studies in NHPs, an alternative anesthetic regimen that resembles the awake state will be optimal. In rodents, continuous infusion of the selective α2-adrenergic agonist medetomidine has been shown to provide analgesia and sedation for several hours while preserving neurovascular coupling, which is vital for detecting both task-based activations and resting-state fluctuations (Kalthoff et al., 2013; Pawela et al., 2009; Sirmpilatze et al., 2019; Weber et al., 2006). Medetomidine - or its active isomer dexmedetomidine - exerts its hypnotic and sedative effect mainly by activating the α2-adrenergic receptors in the locus coeruleus (Sinclair, 2003). It also agonizes the activity of imidazoline type 2 (I2) receptors, which may be one pathway of its neuroprotective effects (Kaur and Singh, 2011). On the negative side, medetomidine also induces hemodynamic changes, with the most prominent being reflex bradycardia due to hypertension caused by initial vasoconstriction (Bol CJJG et al., 1997; Sinclair, 2003). Supplementing medetomidine infusion with low concentrations of isoflurane is used to counterbalance its strong bradycardic effects and to preserve a more favorable physiological condition (Brynildsen et al., 2017). Previously we showed that most of the metabolic effects observed under isoflurane (e.g. increase in brain lactate and myo-inositol) were attenuated by medetomidine (Boretius et al., 2013), further supporting the combined use of medetomidine with isoflurane. Furthermore, this combination protocol, hereafter referred to as med-ISO, represents the current consensus choice for rodent resting-state fMRI studies due to its ability to generate biologically plausible RSNs in mice (Grandjean et al., 2020) and rats (Grandjean et al., 2023), with functional network properties similar to the awake state (Grandjean et al., 2023, 2020).

Despite these promising results, the efficacy of the med-ISO protocol for mapping RSNs in NHPs has not been evaluated in-depth yet and its use only applied sporadically (Ikeda et al., 2023; Ose et al., 2022). Here we adapted the above rodent protocol for acquiring fMRI data in marmoset monkeys and assessed its suitability for task-based and resting-state experiments. Our marmoset med-ISO protocol involved a continuous medetomidine infusion (i.v. 0.1 mg/kg/h) and a low isoflurane concentration (0.4 and 0.6%) administered via a respiratory mask. We began our experiments by examining the concentration-dependent effects of isoflurane-only anesthesia (ISO-only, 1.1 and 1.4%) on visually-evoked BOLD responses compared to med-ISO. We found that BOLD responses were suppressed under ISO-only anesthesia, while they were robust under the med-ISO protocol. Subsequently, we investigated the efficacy of the med-ISO protocol for mapping RSNs through independent-component analysis (ICA) and compared these findings with networks obtained under ISO-only anesthesia or in the awake state (Schaeffer et al. 2022). Lastly, we employed graph-theoretic measures to compare functional connectivity across the awake, med-ISO, and ISO-only states. Our results showed that functional networks under the med-ISO protocol were organized similarly to the awake state, an architecture that was lost under ISO-only conditions.

## Results

### Different brain states under isoflurane-only anesthesia

In primates, isoflurane is routinely used to acquire resting-state fMRI data. However, isoflurane may induce different brain states depending on its concentration and the physiological conditions of each individual. Lower isoflurane concentrations usually induce high-amplitude low-frequency activity, often referred to as the slow-wave state. With increasing isoflurane concentration, the brain reaches the burst-suppression state, in which bursts of activity are interleaved with periods of quiescence (suppressions). At even higher isoflurane concentrations, the quiescent periods are progressively prolonged, leading to a persistent suppression of cortical activity (Kroeger et al., 2013; Kroeger and Amzica, 2007). In recent work (Sirmpilatze et al., 2022) we showed that burst-suppression could be reliably identified in fMRI data using an approach based on principal component analysis (PCA) and that this state underlies the cortico-striatal synchronization observed by multiple fMRI studies in isoflurane-anesthetized subjects (Golkowski et al., 2017; Kalthoff et al., 2013; Liu et al., 2013; Paasonen et al., 2018; Zhang et al., 2019). Nevertheless, burst-suppression only appears in a subset of fMRI runs (Sirmpilatze et al., 2022), meaning the rest likely correspond to other isoflurane-induced brain states. Given these results, we hypothesized that the different states would be reflected on the overall network pattern.

To investigate this possibility, we adapted our PCA-based approach for detecting burst-suppression and applied it to a novel whole-brain fMRI dataset acquired in 8 common marmosets (3 females) at 9.4 Tesla. Multiple RS-fMRI runs were obtained in each monkey, covering a wide range of isoflurane concentrations (1.1 - 1.7%) and summing up to 370 min in total. We extracted the BOLD signal time series from cortical and striatal voxels and visualized them as a carpet plot - i.e. a heatmap of BOLD signal intensity across time (**Figure 1 A**). With this approach we observed multiple distinct events of synchronous signal increase, which were captured by the first temporal principal component (PC1, **Figure 1 C**). PC1 was strongly correlated with the striatum and most of the cortex but spared visual cortices (**Figure 1 B**). As we have shown, this spatial distribution corresponds to a map of burst-suppression, hence we will hereafter refer to it as ‘burst-suppression network’ (Sirmpilatze et al., 2022). We have identified PC1 as the direct hemodynamic correlate of burst-suppression activity (Sirmpilatze et al., 2022), with its peaks corresponding to bursts.

**Figure 1:**
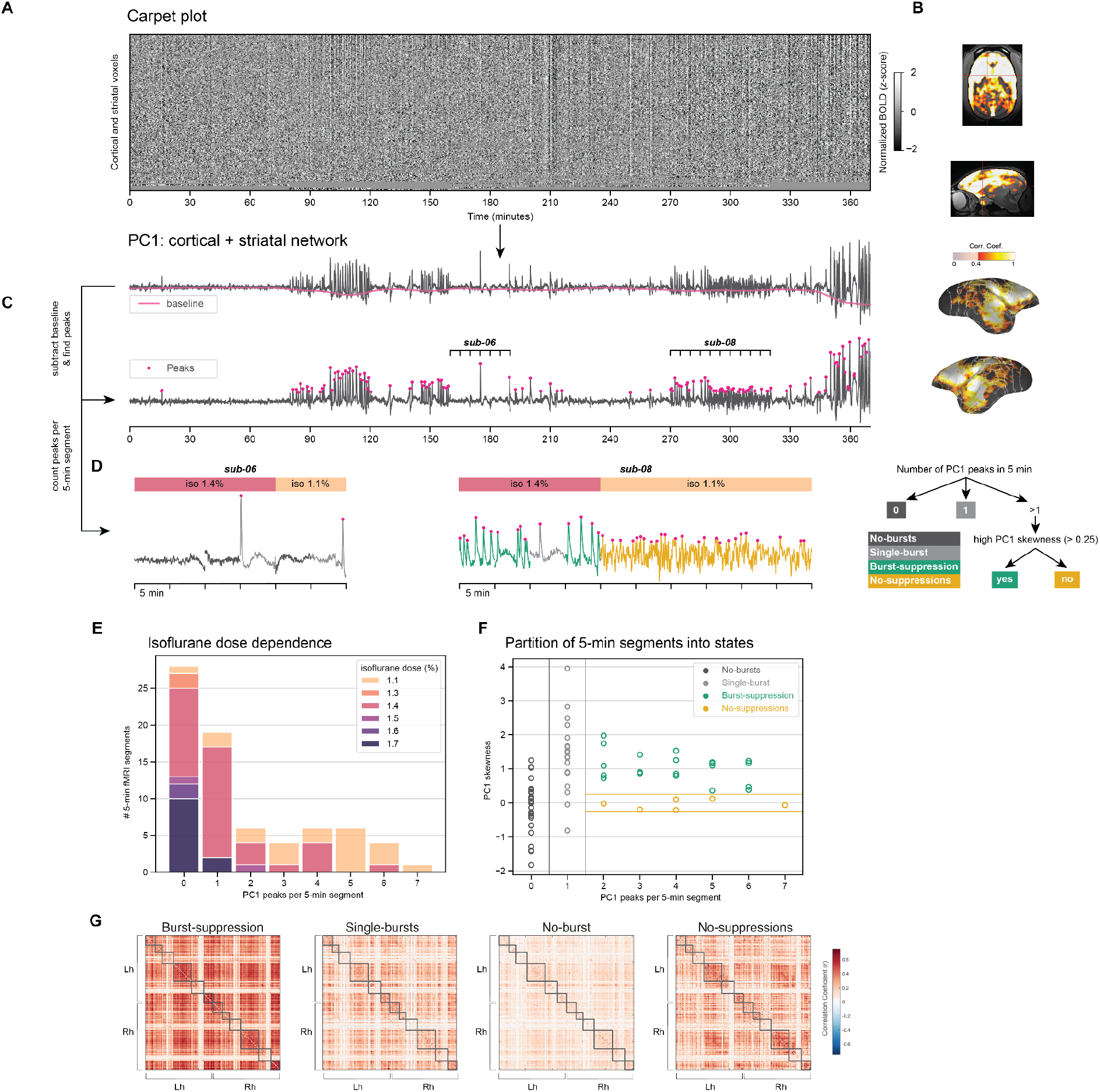
Different brain states under isoflurane anesthesia. **A**. Carpet plot of the blood-oxygen-level-dependent (BOLD) signal across the cortex and the striatum, derived from resting-state (RS) fMRI runs acquired with isoflurane-only (ISO-only) anesthesia. Each row shows the normalized (z-scored) BOLD signal of a single voxel, across all RS fMRI runs. The rows are ordered according to their correlation with the mean across-voxels signal (decreasing from top to bottom). **B**. The first temporal principal component (PC1) of the carpet matrix is strongly correlated with the striatum and most of the neocortex but spares visual cortices (“burst-suppression network”). The Pearson correlation map (thresholded at >0.4) is plotted on the marmoset template brain. **C**. The PC1 time series exhibits sharp peaks - presumed to be associated with bursts. Burst peaks are detected based on the detrended PC1, and the number of peaks is counted per 5-min segment of the time series. **D**. Examples of two subjects with relatively sparse (sub-06) and dense (sub-08) peaks are highlighted. The PC1 time series of sub-08 during the isoflurane concentration of 1.1%, unlike the rest of the dataset, did not exhibit clear peaks. **E**. A histogram of the number of peaks per 5-min segment is shown for all 74 segments, colored according to the isoflurane concentration during the segment’s acquisition. Lower isoflurane concentrations are associated with denser peaks. **F**. The 5-min fMRI segments are partitioned into states based on the number of peaks - 0 (no bursts), 1 (single burst), or >2 (burst-suppression). The plot shows the number of peaks per 5-min segment against the skewness of the PC1 signal within the corresponding segment. The aforementioned sub-08 segments acquired at 1.1% exhibit near-zero skewness and were classified as a separate state (no suppressions). **G.** Functional connectivity matrices for the four partitioned states. Gray boxes indicate modules (n = 6 per hemisphere) detected based on the burst-suppression state connectivity matrix.

The density of PC1 peaks (bursts) varied considerably across the 370 min of recordings, including some extended periods without peaks. We split the data into non-overlapping 5-minute segments and counted the number of burst peaks per segment (**Figure 1 C - D**). 35 % (28/74) of segments showed no peaks, while segments with denser peaks occurred at lower isoflurane concentrations (1.1 - 1.4%, **Figure 1 E**). We partitioned the segments into the following states: ‘no bursts’ (0 peaks), ‘single burst’ (1 peak), and ‘burst-suppression’ (≥ 2 peaks). In one subject (sub-08) we detected burst-suppression during segments acquired with isoflurane 1.4%, but the clear separation between peaks was lost at 1.1%. We attributed this to a lack of suppression periods and assigned the corresponding 6 segments to a fourth ‘no suppressions’ state. The lack of clear peak separation was also reflected in the near-zero skewness of these segments’ time series, whereas most burst-suppression segments showed a highly positive skew (**Figure 1 F**).

Next we grouped the 74 fMRI segments into the four aforementioned states and calculated each state’s functional connectivity matrix. The functional connectivity profiles differed across states, with burst-suppression showing a four-fold increase in edge strength in comparison to the no-bursts state which exhibited globally suppressed connectivity values (**Figure 1 G** and **supplementary figure 1**). This variety in isoflurane-induced brain states and their corresponding effects on functional connectivity may lead to contradictory results within and between studies. Moreover, since burst-suppression and persistent suppression (i.e. no bursts) radically differ from the awake state in neurophysiological terms, one may question the validity of comparisons between isoflurane-anesthetized and awake fMRI studies.

To overcome these confounding effects of isoflurane anesthesia, we decided to investigate an alternative anesthetic protocol that relies on the constant infusion of the sedative agent medetomidine, supplemented with small amounts of isoflurane delivered via a mask. As mentioned above, this med-ISO protocol currently represents the consensus choice for fMRI studies in rodents (Grandjean et al., 2023, 2020) but its effectiveness in nonhuman primates remains unknown. In the following sections, we systematically compare the med-ISO protocol (medetomidine supplemented with either 0.4% or 0.6% of isoflurane) with ISO-only anesthesia and with the awake state.

### Suppressed visual responses under isoflurane-only anesthesia

First we examined the strength of visually-evoked BOLD responses during ISO-only anesthesia at concentrations of 1.1% and 1.4%. Five marmosets (N = 5 subjects) were exposed to visual stimulation via an LED flickering light. At the group level, we observed only weak activations along the visual pathway (see group map of *B*-coefficients in **Figure 2 A**). Significant voxels were mapped based on a Nearest-neighbor cluster with a minimum size of 50 voxels and a level of significance of p < 0.05 (uncorrected) and a Z-score > 2. Beta coefficients ranged between −0.2 to 0.28, Z-scores ranged between −4.3 to 4.1, and FDR q < 0.9. Significant activation was observed in 5 central clusters, including left and right V1, left and right pulvinar, and right LGN (see cluster results in **Table 1**). We believe that this asymmetry in LGN was observed by chance and does not relate to a physiological or anatomical bias (see also **Supplementary Figure 2**). The activation clusters in V1 were mainly concentrated along the medial wall of V1 and spared most of the lateral and foveal areas. No activation was found in area MT+.

**Figure 2:**
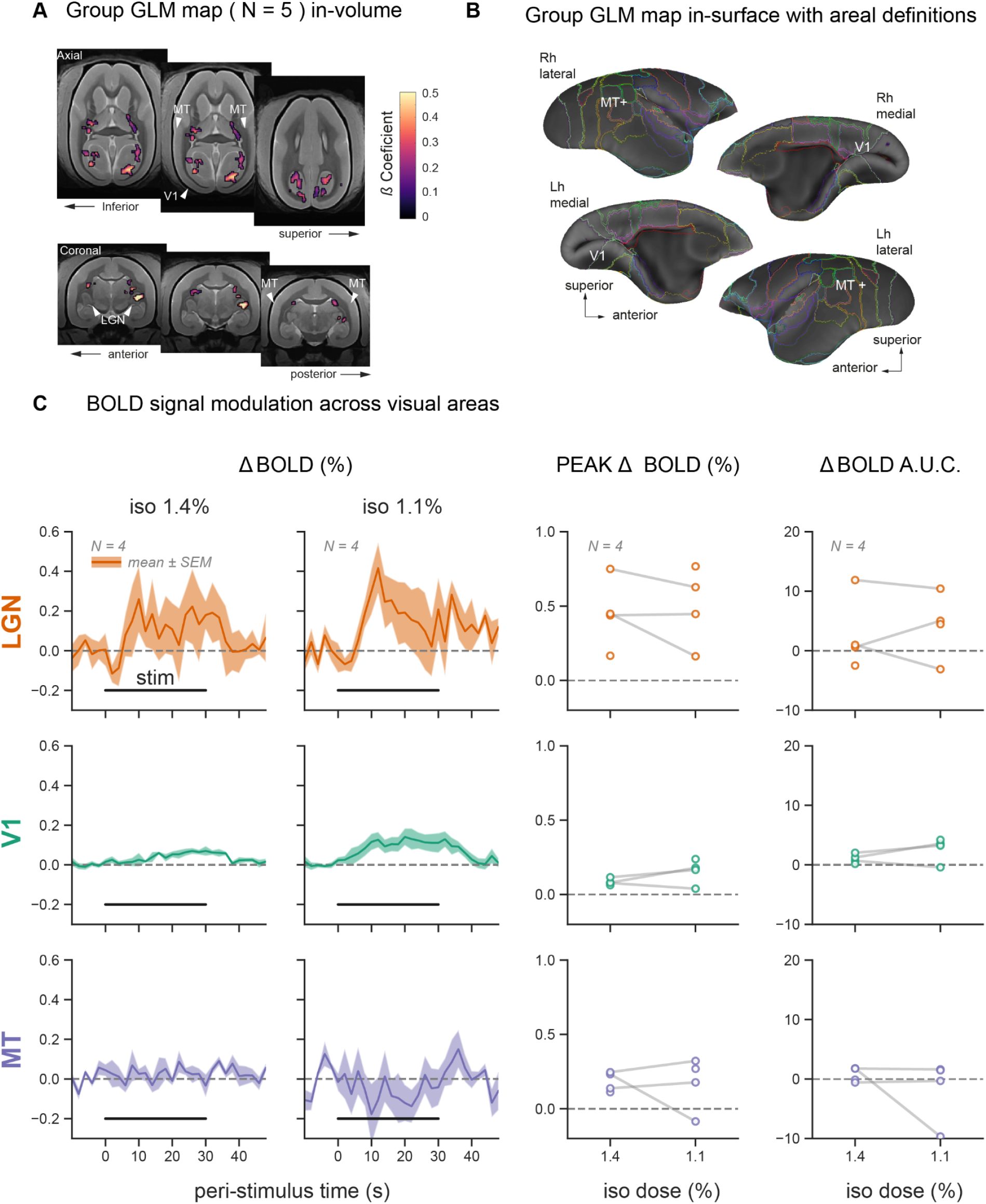
Evoked visual responses are suppressed under isoflurane-only (ISO-only) anesthesia. **A**. During visual stimulation under ISO-only anesthesia, we observed a weak functional activation in the visual system on the group level (N = 5 subjects). The maps are based on a t-test across subjects. Significant voxels were mapped based on a cluster size of 50 voxels and set at a p-value < 0.01. Significant but weak activation was observed in the lateral geniculate nucleus (LGN, only unilaterally) and parts of the medial primary visual cortex (V1). Higher-level motion temporal area MT+ was not activated. **B**. Same group activation map plotted on the cortical surface further confirms a lack of higher-level activation in area MT+. **C**. Responses to visual stimulation are shown as % BOLD signal change in three regions of interest: LGN, V1, and visual area MT. Visual stimulation blocks were averaged within each subject, separately for the two isoflurane levels—1.4% and 1.1%. The mean ± SEM response traces across subjects (N = 5) are plotted on the left. The peak % BOLD response and the area under the curve (AUC) were extracted from each subject’s average trace during stimulation (0 – 30 s). These metrics are plotted across the two isoflurane levels on the right (within-subject measurements are connected via gray lines).

**Table 1:**
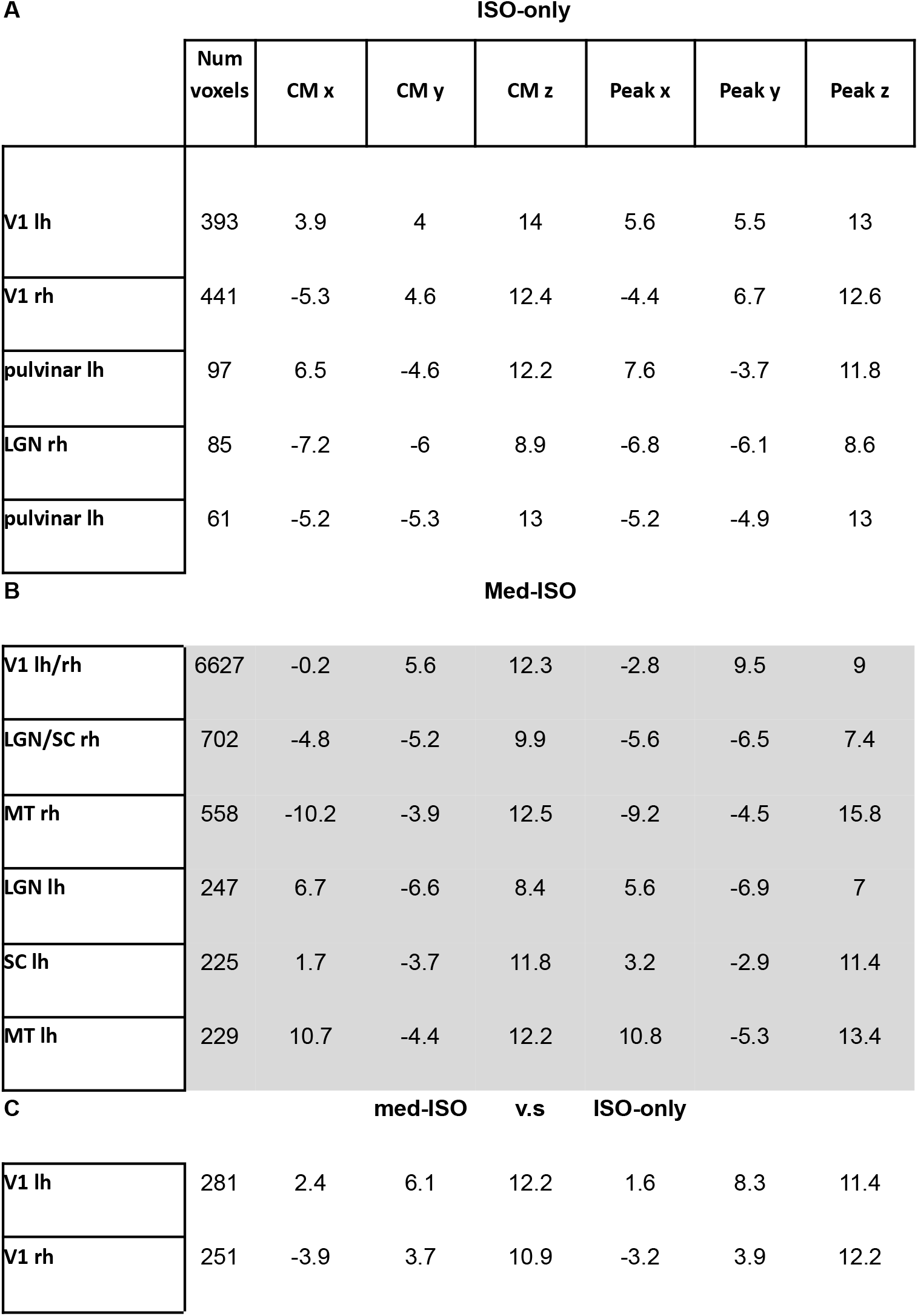
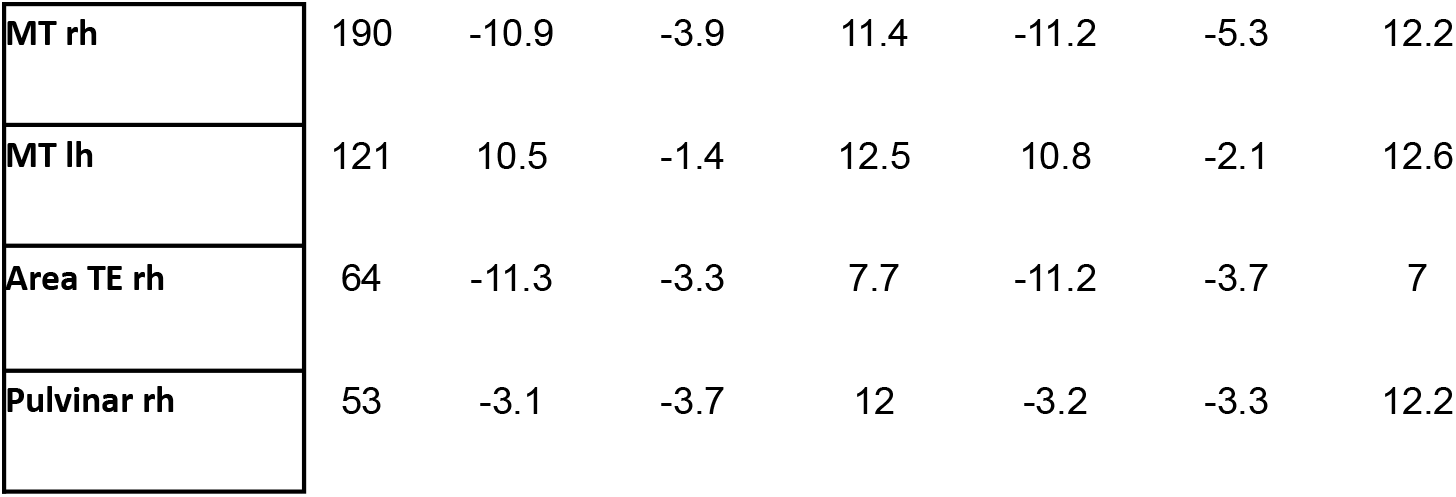
Significantly activated regions showing their cluster size, area label, and area coordinate. The coordinates highlight either the center of mass (CM) or the peak cluster in RAI coordinates from the atlas. The activation regions also show the number of voxels for that cluster. All data were thresholded at a significant p-value of < 0.05, a T-value > 2, and at a cluster minimum size of 50 voxels. **A**. Shows the significant active regions for med-ISO conditions. **B**. Shows the significant active regions for ISO-only conditions. **C**. Shows the significant active regions for the contrast between med-ISO and ISO-only conditions.

**ISO-only**
|  | Num voxels | CM x | CM y | CM z | Peak x | Peak y | Peak z |
| --- | --- | --- | --- | --- | --- | --- | --- |
| V1 lh | 393 | 3.9 | 4 | 14 | 5.6 | 5.5 | 13 |
| V1 rh | 441 | -5.3 | 4.6 | 12.4 | -4.4 | 6.7 | 12.6 |
| pulvinar lh | 97 | 6.5 | -4.6 | 12.2 | 7.6 | -3.7 | 11.8 |
| LGN rh | 85 | -7.2 | -6 | 8.9 | -6.8 | -6.1 | 8.6 |
| pulvinar lh | 61 | -5.2 | -5.3 | 13 | -5.2 | -4.9 | 13 |
| **Med-ISO** |  |  |  |  |  |  |  |
| V1 lh/rh | 6627 | -0.2 | 5.6 | 12.3 | -2.8 | 9.5 | 9 |
| LGN/SC rh | 702 | -4.8 | -5.2 | 9.9 | -5.6 | -6.5 | 7.4 |
| MT rh | 558 | -10.2 | -3.9 | 12.5 | -9.2 | -4.5 | 15.8 |
| LGN lh | 247 | 6.7 | -6.6 | 8.4 | 5.6 | -6.9 | 7 |
| SC lh | 225 | 1.7 | -3.7 | 11.8 | 3.2 | -2.9 | 11.4 |
| MT lh | 229 | 10.7 | -4.4 | 12.2 | 10.8 | -5.3 | 13.4 |

**Table 1:** Significantly activated regions showing their cluster size, area label, and area coordinate.
|  |  |  |  |  |  |  |  |
| --- | --- | --- | --- | --- | --- | --- | --- |
| V1 lh | 281 | 2.4 | 6.1 | 12.2 | 1.6 | 8.3 | 11.4 |
| V1 rh | 251 | -3.9 | 3.7 | 10.9 | -3.2 | 3.9 | 12.2 |
| MT rh | 190 | -10.9 | -3.9 | 11.4 | -11.2 | -5.3 | 12.2 |
| MT lh | 121 | 10.5 | -1.4 | 12.5 | 10.8 | -2.1 | 12.6 |
| Area TE rh | 64 | -11.3 | -3.3 | 7.7 | -11.2 | -3.7 | 7 |
| Pulvinar rh | 53 | -3.1 | -3.7 | 12 | -3.2 | -3.3 | 12.2 |

Next, we extracted BOLD response time series from three main visual brain areas, the LGN, V1, and area MT+ for each individual subject. Peri-stimulus epochs from −10 sec to +50 sec relative to stimulus onset were extracted, normalized as % signal change relative to the pre-stimulus baseline (△BOLD(%)), and averaged within subjects for each isoflurane level. We observed comparable BOLD responses for ISO-only concentrations of 1.4% and 1.1% in the LGN. A weak BOLD response was found in V1 with a slightly stronger signal change at the lower ISO concentration. At both ISO concentrations, no evoked BOLD response was observed in area MT+ (see **Figure 2 C**). To further quantify the concentration-dependent effects of ISO, we calculated the peak and the area under the curve (AUC) of △BOLD(%) and plotted them separately for each concentration (see right panels of **Figure 2 C**). No statistical comparison was performed between concentrations, because paired measurements were only available in 3/5 marmosets.

Having observed weak stimulus-driven activation of the visual system under the ISO-only protocol, we next evaluated whether we would get stronger responses in the visual system with med-ISO anesthesia.

### Robust visual responses under med-ISO anesthesia

We repeated the same experiments under med-ISO anesthesia and evaluated the significance of the visually driven BOLD response in the same cohort of subjects (N = 8). We observed robust BOLD responses to visual flicker stimulation in each individual subject. Most active clusters could be seen along the visual pathway and included bilateral activation of the lateral geniculate nucleus (LGN), bilateral activation of the primary visual cortex (V1), bilateral activation of area MT+ and, interestingly, bilateral activation of the superior colliculus (SC). As shown exemplarily for one monkey in **Supplemental Figure 3**, under med-ISO, the flickering light paradigm effectively evoked BOLD responses even though the eyelids were closed and the eyes were covered with eye creme. The observed BOLD responses in V1 robustly followed the presentation rate of stimulation, and significant BOLD responses could be observed in all respective regions of the pathway.

We found significant activation along the visual pathway also at the group level (see **Figure 3 A**). The group map was obtained based on each subject’s *B*-coefficient estimates. Significant voxels were mapped based on a Nearest neighbor cluster with a minimum size of 50 voxels, a significance level of p < 0.05 (uncorrected) and a Z-score of > 2. Beta coefficients ranged between −0.5 to 0.19, Z-score ranged between −4 to 4.6, and FDR q < 0.5. Significant activation was observed in 6 central clusters, including bilateral V1, left and right LGN, left and right MT+ area, and left and right SC (see cluster results in **Table 1**).

**Figure 3:**
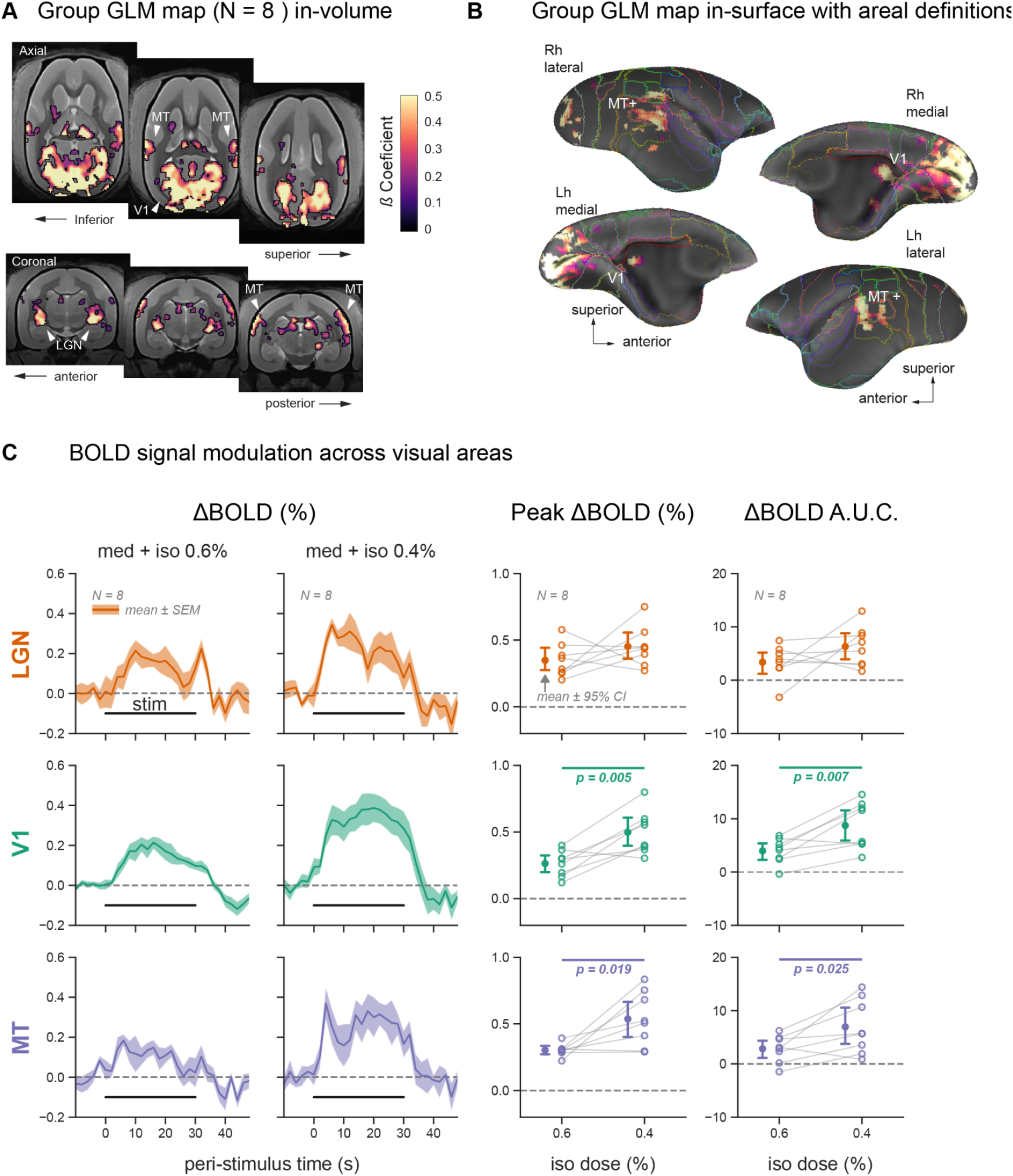
Robust visual responses under medetomidine and low isoflurane (med-ISO) anesthesia. **A**. During visual stimulation under med-ISO anesthesia, we observed significant functional activation in the visual system at the group level (N = 5 subjects). The activation map was based on a t-test across the activation maps obtained in each subject. Significant voxels were mapped based on a cluster size of 50 voxels and set at a p-value < 0.01. Significant activation was observed in visual regions: the lateral geniculate nucleus (LGN), medial primary visual cortex (V1), and motion-sensitive region (MT+). **B**. Same group activation map plotted on the cortical surface. **C**. Responses to visual stimulation are shown as % BOLD signal change in three regions of interest: LGN, V1, and visual area MT. Visual stimulation blocks were averaged within each subject, separately for the two isoflurane levels—0.6% and 0.4%. The mean ± SEM response traces across subjects (N = 8) are plotted on the left. The peak % BOLD response and the area under the curve (AUC) were extracted from each subject’s average trace during stimulation (0 – 30 s). These two metrics were compared across the two isoflurane levels via paired T-tests (p-values indicated when p < 0.05, error bars represent mean ± 95% confidence intervals across subjects, and within-subject measurements are connected via gray lines).

To evaluate the extent and reliability of functional activation along the visuo-geniculate pathway, we calculated the degree of overlap between the activation maps and ROIs obtained from the atlas parcellation. We confirmed an overlap between the activation maps and the ROIs in the LGN and area MT+ (see **Supplementary Figure 4 A**). Mapping the activation on the cortical surface also confirmed the activation of the superior temporal area MT+ (**Figure 3 B**). The activation in area V1 was mainly concentrated along the medial wall of V1 and spared most of foveal V1.

Next, we analyzed BOLD responses from the three brain areas robustly activated by visual stimulation during med-ISO anesthesia (LGN, V1, and area MT+) in each individual subject. Visual stimulation blocks were averaged within each subject, separately for both med-ISO conditions (med-ISO 0.6% and 0.4%). We evaluated △BOLD(%) in the three regions to explore the potential effect of the different isoflurane concentrations. We observed a robust BOLD response across all three ROIs (see **Figure 3 C**) for both med-ISO conditions, with the lowest signal change in area MT+. Compared to med-ISO 0.6 %, an increased BOLD response was observed for med-ISO 0.4 %, with the highest △BOLD(%) in V1. The overall response difference between the two med-ISO conditions was 0.2%. We calculated the peak BOLD response (Peak △BOLD (%) and the area under the curve (△BOLD(%) A.U.C) to further quantify these concentration-dependent effects of isoflurane (see right panels of **Figure 2 C**). Both response metrics showed a significant difference between the two med-ISO conditions as confirmed by pairwise comparison across the subjects (N = 8, paired t-test, p-value < 0.05).

Given the observed robust BOLD response under med-ISO and the weak change in BOLD signal under ISO-only, we next compared the two anesthesia regimes directly, using the 5 subjects in which visual stimulation data was acquired with both protocols. For this comparison, we pooled the two med-ISO conditions (med-ISO at 0.4% and 0.6%) and the two ISO-only conditions (ISO-only at 1.1% and 1.4%), respectively.

### Stronger visual activation under med-ISO compared to ISO-only anesthesia

We analyzed the same group of marmoset subjects (N = 5 subjects) to compare the difference between the two anesthetic conditions, med-ISO versus ISO-only. We observed a significantly stronger activation along the visuo-geniculate pathway for med-ISO (yellow-red) in contrast to the ISO-only condition (cyan-blue, see **Figure 4 A**). The contrast map was obtained based on a paired t-test between *B*-coefficients of each condition for the same subject. Significant voxels were mapped based on a nearest neighbor cluster with a minimum size of 50 voxels, a level of significance p < 0.05 (uncorrected) and a Z-score > 2. Beta coefficients ranged between −0.42 to 0.36, Z-score ranged between −4.7 to 4.3, and FDR < q 0.9. Significant activation was observed in 6 central clusters, including left and right V1, left and right MT, right area TE and right pulvinar (see cluster results in **Table 1**). In V1, the largest effect size was mainly observed for med-ISO conditions along the medial wall of both hemispheres. A similar contrast difference was observed for area MT+, where the med-ISO condition elicited a more significant response in these higher-level regions. Mapping the contrast results on the cortical surface further confirmed the stronger visual activation of area MT+ and medial V1 (see **Figure 4 B**) under med-ISO. At the level of the LGN, we notice no contrast difference for ISO-only or med-ISO. Next, we directly compared the BOLD responses from the LGN, V1, and area MT+ obtained under med-ISO and ISO-only. For both conditions, we observed a BOLD response at the level of the LGN. However, only under the med-ISO condition the BOLD response could be driven beyond the thalamus towards V1 and the higher-level motion-sensitive area MT+ (see **Figure 4 C**, same ROIs as in **Figure 2 C** and **3 C**).

**Figure 4:**
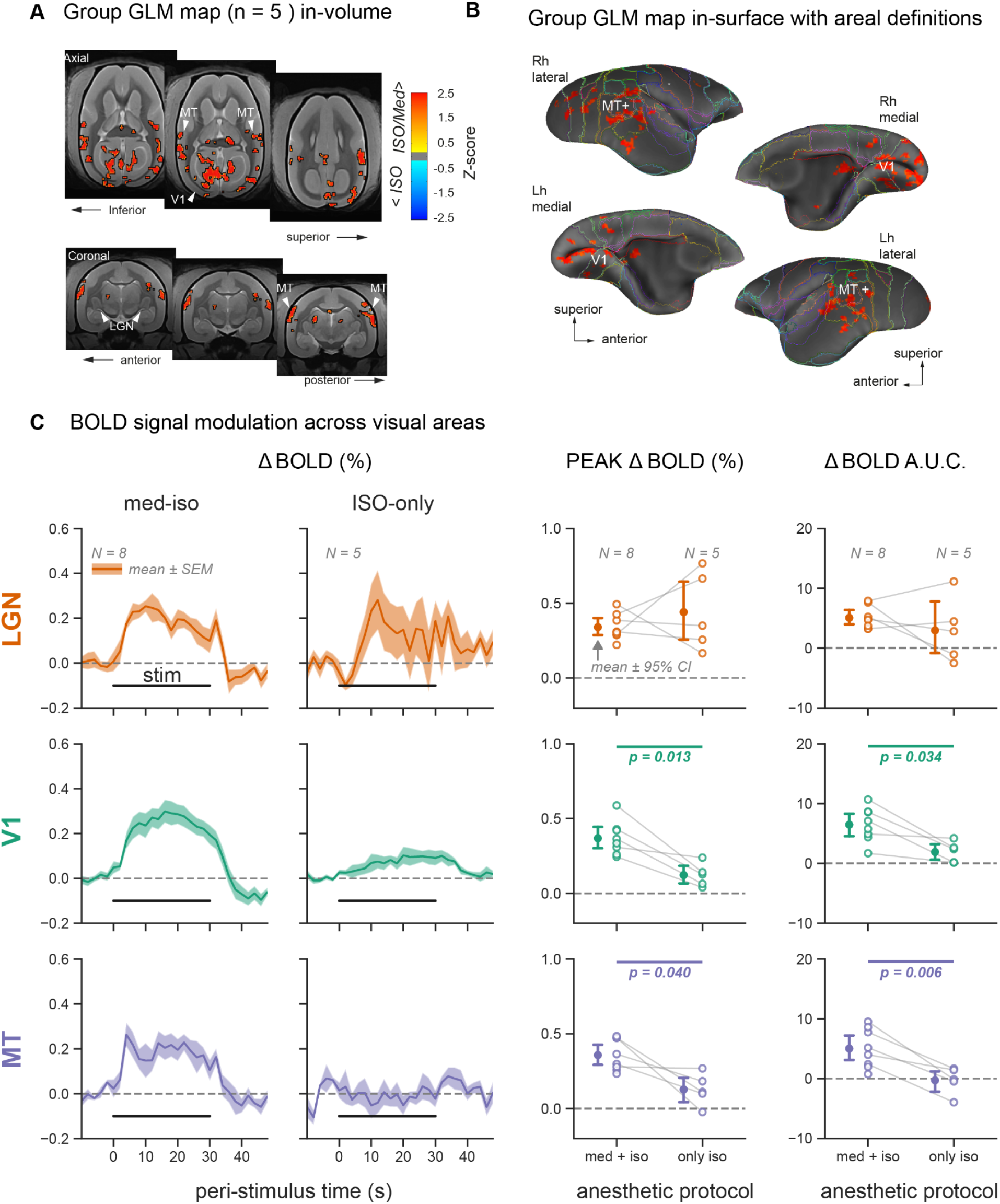
Contrast between anesthetic regimens reveals higher sensitivity in visual areas for med-ISO compared to ISO-only. **A**. Pairwise group level (N = 5 subjects) contrast map between med-ISO and ISO-only anesthesia. The contrast shows a greater response for med-ISO in the visual system. Significant voxels were mapped based on a cluster size of 50 voxels and set at a p-value < 0.01. Contrast was observed in the left lateral geniculate nucleus (LGN), in the medial primary visual cortex (V1) bilaterally and in the higher-level motion temporal area MT+ bilaterally. **B**. Same contrast map plotted on the cortical surface further confirms the stronger activation of areas V1 and MT+ for med-ISO anesthesia. **C**. Responses to visual stimulation are shown as % BOLD signal change in three regions of interest: LGN, V1, and visual area MT. Visual stimulation blocks were averaged within each subject, separately for the two anesthetic protocols: med-ISO and ISO-only. The mean ± SEM response traces across subjects (N = 8 for med-ISO and N = 5 for ISO-only) are plotted on the left. The peak % BOLD response and the area under the curve (AUC) were extracted from each subject’s average trace during stimulation (0 – 30 s). These two metrics were compared across the two anesthetic regimens via paired T-tests (p-values indicated when p < 0.05, error bars represent mean ± 95% confidence intervals across subjects, and within-subject measurements are connected via gray lines).

The response metrics, peak △BOLD (%) and △BOLD A.U.C. showed a considerable difference between the two anesthesia conditions in both cortical area V1 and area MT+ but not for the LGN. A pairwise T-test between the two anesthesia protocols confirmed a significant difference between the two conditions (N = 5, p-value < 0.05) in the two cortical regions.

Overall, these analyses highlight the effectiveness of the med-ISO anesthesia protocol in driving the visual system’s BOLD response in marmosets compared to ISO-only anesthesia. Having evaluated the effectiveness of the med-ISO protocol in driving the BOLD response, we next explore the degree to which the med-ISO protocol could be used to detect common RSNs in marmosets compared to ISO-only and awake conditions.

### Mapping common RSNs under different anesthesia conditions using ICA

In addition to task-based-fMRI for driving the BOLD response in the visual system, we were also interested in evaluating the degree to which RSNs remain present under the different anesthetic states. To evaluate this, we also obtained resting-state data during the same imaging session. We conducted these experiments for med-ISO and ISO-only to evaluate how anesthesia conditions affected the RSNs commonly observed in the awake marmoset.

The RSNs identified by group-based independent component analysis (ICA) of the med-ISO data are shown in **Figure 5**. We evaluated the networks based on visual inspection and with the aid of the well-described resting-state brain patterns in awake marmosets. Inspection criteria included symmetrical patterns, non brain edge patterns, or scatter clusters. For each identified network, we also report its explained variance (e.v.) and total variance (t.v.). A threshold of p-value <0.05 and a minimum Z-score of 2 were used for all networks. All maps were clipped at a maximum Z-score value of (+/−10). Using these criteria, we identified the following networks: The pre-motor network (4.86 % e.v.; 1.69 % t.v.), the default-mode network (4.66 % e.v.; 1.63 % t.v.), the sensory-motor network (4.40 % e.v.; 1.54 % t.v.), the basal ganglia network (3.93 % e.v.; 1.37 % t.v.), the visual network I (3.88 % e.v.; 1.35 % t.v.) and visual network II (Attention, 3.86 % e.v.; 1.34 % t.v.). Performing the same analyses on the openly available awake marmoset’s resting-state data, we identified similar RSNs (see **Supplementary Figure 5 and 6**). On the other hand, performing the same analyses on resting-state data from the same subjects under the ISO-only condition resulted in dramatically different components, with the cortical and striatal parts of the burst-suppression network as the predominant components (see **Supplementary Figure 7**).

**Figure 5:**
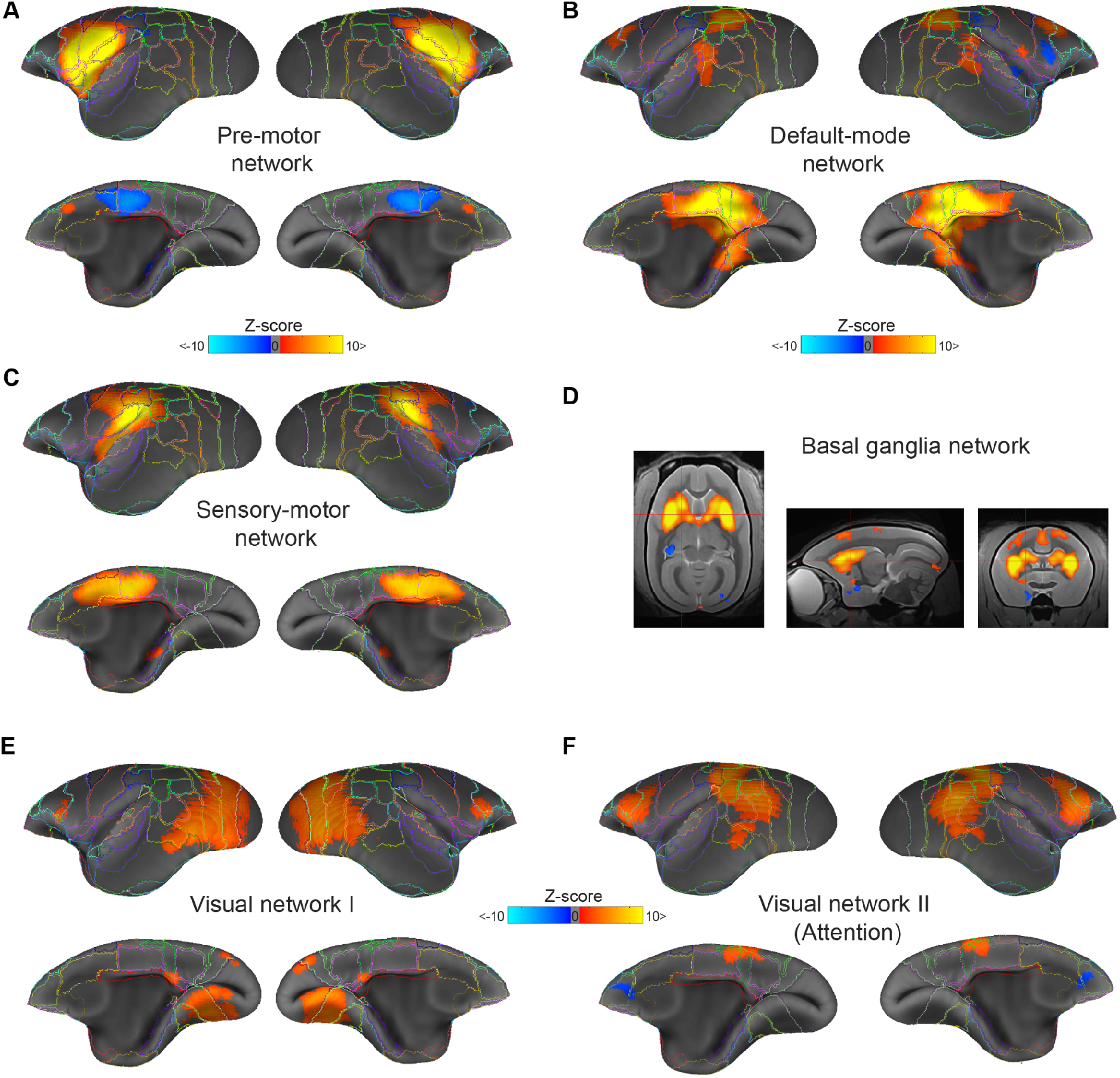
Robust and reliable resting state networks obtained under medetomidine and low isoflurane (med-ISO) anesthesia. Group-level independent component analyses identified commonly reported resting-state networks in the common marmoset under med-ISO anesthesia. All networks were thresholded at a p-value < 0.05 and a minimum Z-score of 2 and clipped at a maximum Z-score value of (+/−10). Using these criteria, we identified the following networks: (**A**) the pre-motor network (4.86 % e.v.; 1.69 % t.v.), (**B**) Default-mode network (4.66 % e.v.; 1.63 % t.v.), (**C**) the sensory-motor network (4.40 % e.v.; 1.54 % t.v.), (**D**) the basal ganglia network (3.93 % e.v.; 1.37 % t.v.), (**E**) the visual network I (3.88 % e.v.; 1.35 % t.v.) and (**F**) visual network II (Attention, 3.86 % e.v.; 1.34 % t.v.). See supplementary figure 4 and **5** for the identified ICa networks under awake and ISO-only respectively. e.v. - explained variance; t.v. - total variance.

Based on our ICA results, we conclude that med-ISO anesthesia in marmosets resulted in qualitatively similar patterns of RSN activity as those observed in the awake state. In contrast, these RSN patterns strongly differed from those obtained under ISO-only.

ICA allowed us to identify RSNs under different anesthesia protocols and to match them qualitatively. However, directly comparing networks using the ICA approach remains a challenge. Thus, we next used graph-theoretical measures to quantify and evaluate the differences in network structure across anesthesia conditions, including those obtained in the awake state.

### Functional network structure is preserved under med-ISO anesthesia

In contrast to the qualitative comparisons of anesthesia conditions using ICA, graph analyses enabled us to compare RSNs quantitatively. To observe the effects of anesthesia on RSNs, we constructed connectivity matrices for each network condition; awake, med-ISO 0.4%, med-ISO 0.6%, and ISO-only. Since the awake condition represents our ideal, reference condition, we compared changes in network connectivity departing from the awake state. Toward this end, we first calculated the community organization of the awake network. Then, with the identified network communities we reorganized all other network states (e.g. med-ISO 0.4%, med-ISO 0.6%, and ISO-only) based on this modularity structure. Modularity was calculated on the average awake connectivity matrix of both cerebral hemispheres of all subjects. On this average matrix, the Louvain community algorithm detected four modules that mostly captured frontal (FC), subcortical (Sub), motor (MC), and visual cortex (VC) (see **Figure 6 A**). Visual inspection of the matrices indicated similar communities among the awake, the med-ISO 0.4, and med-ISO 0.6 matrices but stark difference to the average ISO-only matrix. In previous analysis, we segregated ISO-only matrices into multiple burst states (e.g., burst, mostly burst, and no burst, **Figure 1**). To compare these results with the awake and med-ISO states, we reorganized all matrices according to the four modules per hemisphere we found under the awake condition. Most of the ISO-only states showed a global increase in strength with a complete loss of the modular organization seen in the awake state. In contrast, under med-ISO, we observed an overall decrease in the strength of the weights with increasing isoflurane concentration (e.g., from 0.4 % to 0.6 %). Despite these dose-dependent effects in weight strength, the network structure remained the same across both concentration conditions.

**Figure 6:**
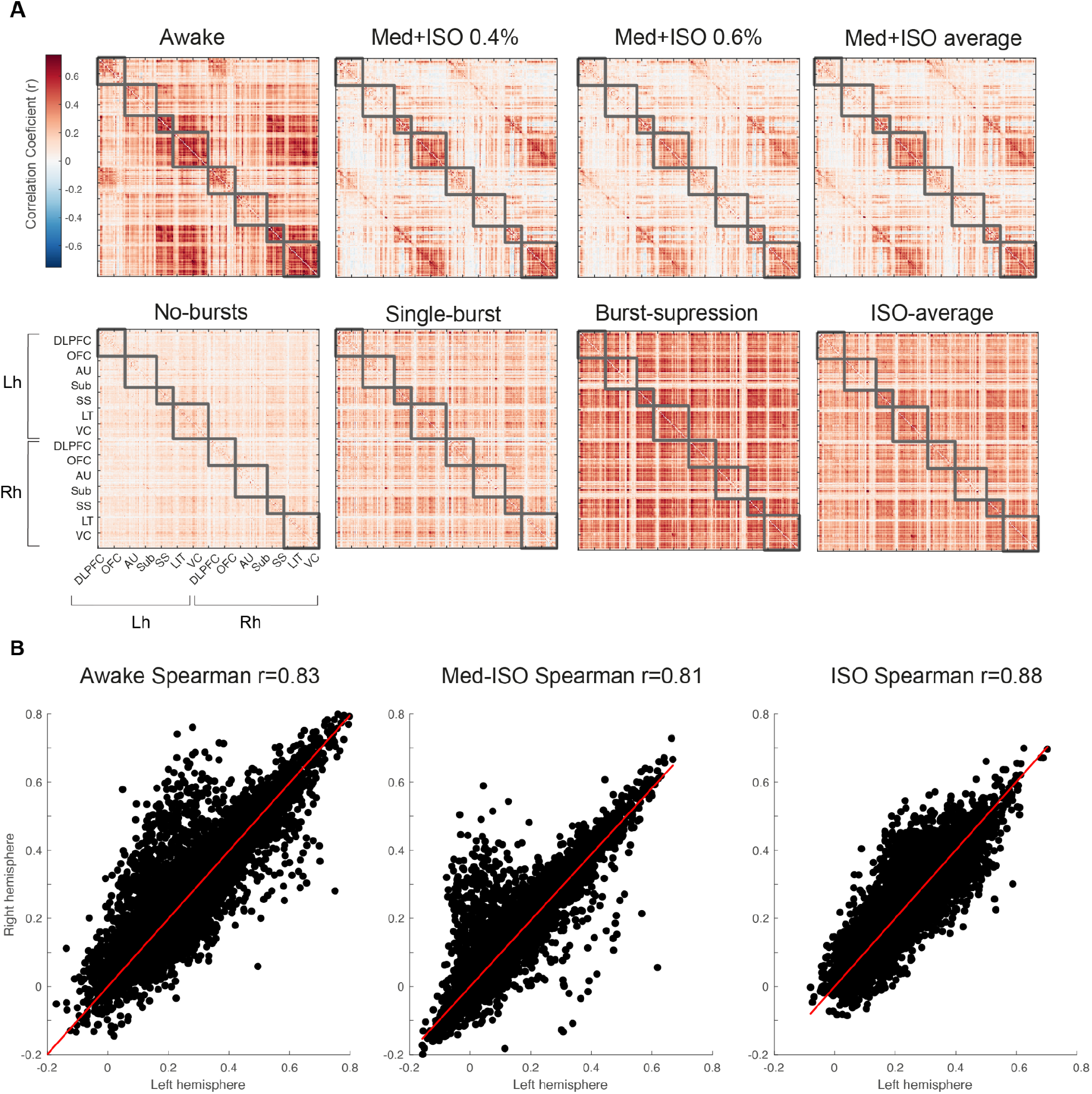
Preservation of functional connectivity structure under med-ISO anesthesia. **A**. Average connectivity matrices for each condition Awake, med-ISO 0.4%, med-ISO 0.6%, and average med-ISO (top). (Bottom) matrices show the ISO-only states for the lack of bursts (no-bursts), the presence of one burst (single-burst), and the clear presence of bursts (burst-suppression), along with the average matrix across all the ISO-only states. The gray squares indicate the maximal no-overlap group based on hierarchical modularity (4 modules per hemisphere) obtained for the awake condition. The Louvain community detection algorithm maximizes the number of within-edges and minimizes the number across groups. The connectivity matrices include each intra-hemispheric correlation and inter-hemispheric correlation. Labels are ordered based on their second-level labeling from the MBV_v3. Labels. The four modules detected largely encompass the frontal cortex (FC), subcortical (Sub), motor cortex (MC), and visual (VC). **B**. Interhemispheric correlation shows the overall connectivity pattern across states; Awake (left), med-ISO (middle), and ISO-only (right). The intersecting line shows the linear fit between the left and right hemispheres. Notice how highly correlated the hemispheres are under the ISO-only condition as compared to the awake and med-ISO conditions.

To compare the modular organization between states we investigated the maximal number of non-overlapping modules per state; awake, med-ISO and those classified from ISO-only conditions, which included burst-suppression, single-bursts and no-bursts. Modularity was calculated separately for each network state. These results showed that the number of module partitions remained the same for awake and med-ISO (n modules = 4) while for the ISO-only conditions the number of modules increased as bursts got sparser (see **supplementary figure 8**).

To further quantify the structure of RSNs across anesthesia conditions, we analyzed the patterns of inter-hemispheric correlations. In general, the patterns appeared similar based on visual inspection of the connectivity matrices. Specifically, the inter-hemispheric correlation coefficients were slightly stronger for ISO-only (spearman’s r = 0.88) and weaker for med-ISO (spearman’s r = 0.76) compared to the awake condition (spearman’s r = 0.82). Moreover, we noted a slight asymmetrical bias for both med-ISO and awake conditions, with some lower-weight connections in the left hemisphere compared to the right hemisphere. This pattern was lost under ISO-only conditions.

We next investigated the connection lengths in RSNs by measuring the connection weights (correlation coefficients) as a function of the Euclidean distance between nodes. Our findings revealed a monotonic decrease in correlation with increasing node distance, consistent with previous human and monkey network studies highlighting the influence of anatomical location and region distance in the brain (Hilgetag and Kaiser, 2004). We distinguished between inter-hemispheric and intra-hemispheric correlations, noting that highly weighted connections typically diminish at longer distances and are more prevalent at shorter distances. This trend was evident in the distribution of connection weights, which peaked at lower weights for both awake and med-ISO conditions but showed a broader distribution with more high-weight instances under ISO-only. This pattern in the ISO-only condition suggests a reduced local segregation and an inefficient network-wide increase in connection strength, deviating from the normal brain connectivity preference for short-range connections to minimize metabolic costs (Shine et al., 2016). Such typical features of brain connectivity, aimed at optimizing the balance between local and wide-range area synchronization, were notably absent in the ISO-only condition.

Finally, comparing intra- and inter-hemispheric connections from the anesthetized conditions (e.g., med-ISO and ISO-only) against the awake condition (**Supplementary Figure 9**), we found a high correlation (adjusted R-square 0.64) between the awake and med-ISO and a relatively lower correlation (adjusted R-square 0.41) between the awake and ISO-only condition.

In summary, under ISO-only, strong correlations were over-emphasized across the whole brain, likely due to the presence of burst-suppression, while our connectivity analysis revealed a similar sub-network architecture for awake and med-ISO conditions. Thus, we conclude that med-ISO is a suitable alternative anesthesia protocol for performing RSN connectivity analyses and task-evoked fMRI in the marmoset monkey.

## Discussion

In this report, we established the use of medetomidine in combination with a low concentration of isoflurane (med-ISO) as an alternative anesthesia protocol for mapping task-based and task-free networks in the marmoset monkey. Our results show that the overall functional network structure is preserved under med-ISO as it closely resembles the awake state. In contrast, our results speak against the use of isoflurane alone as an anesthetic agent for mapping RSNs across species and suggest the combination of med-ISO as an alternative solution for functional brain imaging under anesthesia in non-human primates.

Most resting-state studies on awake - non anesthetized - NHPs converge on a discrete number of spatially distributed patterns that emerge during rest (Tian et al., 2022). To compare these resting state patterns with those found under different anesthetic protocols, we relied on openly available datasets previously acquired in awake marmoset monkeys (Schaeffer et al., 2022). We used this data as a gold standard and processed them similarly to our data.

Using ICA, previous studies identified RSNs in awake marmosets which included the default-mode, pre-motor, sensory-motor, and visual networks, among others (Schaeffer et al., 2022; Tian et al., 2022). These functional networks have been well characterized in both marmosets and macaque monkeys under awake and anesthetized conditions (Hutchison et al., 2014, 2013, 2011). In line with these previous studies, we identified similar functional networks in our med-ISO datasets (**Figure 5**) as those in the awake state (**Supplementary Figure 5 and 6**). However, our findings in ISO-only conditions (**Supplementary Figure 7**) departed starkly from those we obtained under the awake and med-ISO states.

Isoflurane has been extensively used for fMRI studies in NHPs (Hori et al., 2020; Hutchison et al., 2014; Wu et al., 2016; Zhang et al., 2019) with partially contradictory findings. Some studies reported increased functional connectivity (Zhang et al., 2019), while others reported a decrease in functional connectivity (Vincent et al., 2007; Wu et al., 2016). This effect seems to be dose-dependent, where higher concentrations lead to lower cortical activity (Kroeger et al., 2013; Kroeger and Amzica, 2007; Sirmpilatze et al., 2022). Furthermore, under isoflurane, thalamic and hemispheric connectivity were disproportionately and drastically reduced compared to awake conditions (Hutchison et al., 2014). Mixed effects have also been reported in marmosets; some studies indicated the presence of RSNs, while others showed a systemic lack of frontal connectivity under isoflurane (Giacometti et al., 2022; Hori et al., 2020).

Our current results revealed two main independent network components under isoflurane, one emphasizing the basal ganglia and the other comprising the neocortex but sparing visual cortices. The remaining components showed non-specific patterns in the retrosplenial, parietal, and visual cortex. These findings align with our previous results in NHPs and humans (Sirmpilatze et al., 2022), where we showed that within a specific range of isoflurane concentrations (in marmosets, typically between 0.8 and 1.1 %), the brain enters the burst-suppression state; bursts of activity with periods of relative silence that alternate quasi-periodically (Golkowski et al., 2017; Sirmpilatze et al., 2022; Zhang et al., 2019). This state exhibits global synchrony in BOLD signal fluctuations, most marked in the basal ganglia and the neocortex but excluding primary sensory areas, with V1 being the most prominent exception. This burst-suppression network exactly overlaps with the two main independent components we observed here. Interestingly, when we split our current ISO-only dataset based on the appearance of bursts, brain states with no bursts showed significantly lower correlations across the brain. This dependence on anesthesia depth may partially explain the reported contradictions in functional connectivity studies using isoflurane.

Discrepancies in functional connectivity studies may also arise from variations in preprocessing, particularly the use of global signal and motion regression. In our study, we opted not to regress out the global signal. This decision aligns with emerging views that the global signal, rather than being a mere nuisance variable, actually carries valuable information (Liu et al., 2017). This seems especially pertinent in anesthetized states like burst-suppression, where the widespread neural synchrony is likely to be reflected on the global signal. Notably, prior studies have shown that using global signal regression during burst-suppression significantly lowers connectivity values (Kalthoff et al., 2013; Liu et al., 2013; Zhang et al., 2019). A potential workaround could be to average across larger datasets, which might mitigate the confounding effects of isoflurane by diluting them. The necessity of motion regression is debatable for anesthetized measurements, where movement is minimal. We omitted motion regression in our main analysis, as we observed it interfered with detecting bursts in the ISO-only condition (see **Supplementary Figure 10**). However, to ensure the robustness of our findings, we also conducted analyses including motion regression and provided relevant results as supplementary material: **Supplementary figures 5** and **6** illustrate the impact of motion regression on ICA-detected networks in the awake state, while **supplementary figure 11** demonstrates its effect on functional connectivity across states. While we observed some effects, particularly in the med-ISO and awake conditions, motion regression did not significantly change our main findings or conclusions.

As ICA allows for qualitative comparison of network patterns, we next quantitatively explored the functional networks based on the pairwise correlations between the time series of each cortical and subcortical brain region (Ortiz-Rios et al., 2021). We used the awake correlation matrix as our primary network state to extract communities. Here we identified four main modules, largely encompassing frontal, temporal, sensorimotor, and visual areas. We then used the same order of brain regions to create the respective correlation matrices for all anesthesia conditions (**Figure 6 A**). Compared to the awake state, the sub-networks under med-ISO showed a reduced correlation strength which became slightly more apparent with increasing isoflurane concentration. This observation aligns with a previous report in humans where the administration of dexmedetomidine alone significantly diminished the strength of average brain connectivity (Hashmi et al., 2017). However, it is worth noting that, apart from this overall reduction in strength, the network’s modular structure was preserved under med-ISO.

In contrast, the hyperconnectivity observed for ISO-only affected the modular organization and resulted in a loss of the functional network architecture present in the awake state. The increased number of main modules further reflected the loss of network structure under ISO-only (**Supplementary Figure 8**), while for med-ISO the modular organization remained similar to the awake. For the awake and med-ISO conditions, four modules were identified, while the number under ISO-only increased with prolonged periods of suppression. Another characteristic that showed the preserved, awake-like network structure under med-ISO is the slight asymmetry showing more high-weighted connections in the right hemisphere, likewise observable under the awake condition but not under ISO-only (**Figure 6 B**). Interestingly, these asymmetrical patterns arise despite the different source of data acquisitions and state in the openly available data and our med-ISO datasets. In contrast, the network under the ISO-only showed a non-hemisphere specific increase in highly-weighted connections. For the med-ISO instead we observed an increase in negative correlations not seen in the awake state. Our findings are closely consistent with previous rodent studies showing hyperconnectivity in ISO-only conditions due to burst-suppression state and a reduction in connectivity from med-ISO anesthesia (Paasonen et al., 2018). Although, the mechanisms behind the overall decrease in connectivity strength remain to be investigated.

Another important feature of functional connectivity relates to the distance between local or clustered brain nodes which tends to be short. As a consequence, anatomically, nearby regions are considered to be “economical.” In contrast, the edges of long-range projections are fewer and sparse but have the capacity for integration over the network (Watts and Strogatz, 1998). Investigating the correlation strength as a function of node distance, we found a decrease in connection weights as the node distance increased for both med-ISO and awake conditions (**Supplementary Figure 9**). This finding supports the assumption that med-ISO preserves the intrinsic properties of the brain to reduce the metabolic cost of synchronizing activity among a wide range of brain areas by preferring short-range connections. In contrast, under ISO-only high-weighted and low-weighted long distance connections existed to an equal extent indicating a loss of local clustering and a drastic alteration in the macroscale tuning of the network. Moreover, when we compare among the interhemispheric connections we observed a closer correspondence between awake and med-ISO as compared to the awake and ISO-only conditions (**Supplementary Figure 12**) additionally supporting the assumption that med-ISO is a more suitable anesthesia protocol for studying resting-state connectivity.

Considering the number of studies exploring RSNs in NHPs under anesthesia, only a few have investigated the effect of anesthetic protocols on task-based BOLD fMRI. One study in marmosets reported BOLD signal changes under propofol in response to somatosensory stimulation in the thalamus and the primary and secondary somatosensory cortex (Liu et al., 2013). The response amplitudes were significantly attenuated compared to the awake state. To our knowledge, no study in marmosets has yet investigated the effect of anesthesia on the BOLD response to visual stimulation.

Under the proposed med-ISO protocol, we observed robust BOLD responses to visual flicker stimulation on the single subject and group level. Active clusters could be seen along the visuo-geniculate pathway, with strong bilateral activations in LGN, V1, and higher visual cortex MT+. In V1 and MT+, the BOLD response even increased with decreasing isoflurane concentration, as shown by significantly higher peak and A.U.C of △BOLD(%) under med-ISO 0.4% compared to med-ISO 0.6%. This fits well with the reduced cortical activation observed under ISO-only and confirms previous reports on isoflurane-mediated suppression of BOLD responses (Hutchison et al., 2014). Interestingly, this dose dependency was not seen in the LGN. In fact, we found comparable BOLD responses to visual stimulation in the LGN across all anesthetic protocols. Notably, under med-ISO, we also found activations in the superior colliculus (SC), which likely speaks for eye movements in response to the flickering light. Visually-induced BOLD response in the SC is a common observation in rodents under this type of anesthesia (Pradier et al., 2021).

In contrast to the robust BOLD responses along the visual pathway under med-ISO, under ISO-only we found highly variable activations in the LGN and suppressed activations at the level of the primary visual cortex. No task-related responses were seen in area MT+. These findings suggest a disturbed signal propagation along the cortical stream of visual processing, which was preserved under med-ISO. Prior work reported disrupted thalamic activity under isoflurane during resting-state, however, the observed task-based response in LGN indicates that the retino-thalamic drive remained present. Although we did not find activation in higher cortical areas, we detected activation in the pulvinar which may relate to an alteration of reciprocal cortico-thalamic connectivity during visual information processing under isoflurane anesthesia.

Many studies in anesthetized NHPs, avoided the confounding effects of isoflurane by employing various experimental and analytical strategies, such as opting for different anesthetic regimes (Hori et al., 2020; Kalthoff et al., 2013; Logothetis et al., 1999), applying global signal regression during fMRI data preprocessing (Kalthoff et al., 2013; X. Liu et al., 2013; Zhang et al., 2019), or using lower concentrations of isoflurane (X. Liu et al., 2013). Important to consider is the fact that reducing isoflurane concentration requires additional medication to prevent movement and to ensure sufficient anesthetic depth. For instance, in macaques, robust BOLD responses have been reported for 0.2 - 0.5% isoflurane in combination with opiates such as fentanyl or remifentanyl, often with an additional administration of muscle relaxants (Logothetis et al., 2001). That said, using muscle relaxants may be unsuitable from a welfare perspective, especially if the depth of anesthesia can not be adequately monitored. Moreover, the effect of remifentanyl on opioid receptors may be an exclusion criterion for some neuroscience research questions given its addictive properties and speaks against the use of anesthesia protocols that involve the repeated use of opioids, which is of particular interest for follow-up studies with labor-intensive trained NHPs.

With our proposed med-ISO protocol, the isoflurane concentration could be reduced to 0.4 % while maintaining stable anesthesia without the need of additional muscle relaxants. A further reduction of isoflurane led, in our hands, to spontaneous movement in some of the marmoset monkeys, which could not be compensated by increasing the medetomidine dose. Moreover, the infusion of 0.2 mg/Kg/h medetomidine already significantly reduced the heart rate of the monkeys, which we did not pursue further. In contrast, our previous study on rats (Sirmpilatze et al., 2019), consistent with numerous other rodent studies, found that medetomidine infusion alone was sufficient to maintain stable anesthesia for several hours. Thus, we assume this difference might be species-specific.

Another argument for using med-ISO in fMRI studies of anesthetized marmosets is the practicality and ease of use. Isoflurane and oxygen-enriched air can be delivered via a mask in self-breathing animals, eliminating the need for a respirator. The low isoflurane concentration we used did not suppress the breathing rate, as was the case for ISO-only, where intubation was required to ensure sufficient oxygen supply (Pradier et al., 2021). Moreover, medetomidine is reversible by α2 antagonists such as atipamezole, allowing fast and safe recovery of the monkey. Additionally, medetomidine has been shown to have neuroprotective properties and is associated with anti-inflammatory and anti-apoptotic effects. Particularly important in the context of repeated anesthesia in follow-up studies is the fact that it can significantly improve cognition and postoperative outcomes (Kaur and Singh, 2011).

In summary, using ICA, we found similar resting-state networks under med-ISO as those in the awake state but drastically different networks under ISO-only conditions. Task-based fMRI under the med-ISO resulted in robust BOLD responses along the visual pathway up to higher-level regions. Network analysis further confirmed that med-ISO preserves the naturally occurring functional architecture present during resting-state similar to the awake state. The combination of low-isoflurane and medetomidine is a suitable anesthesia protocol that preserves the resting-state network structure and maintains a steady-state of anesthesia ideal for comparative studies based on functional neuroimaging.

## Methods

### Subjects

Imaging experiments were carried out at the German Primate Center (Deutsches Primatenzentrum GmbH, Göttingen, Germany) with the approval from the ethics committee (project number: 33.19-42502-04-17/2496) of the Lower Saxony State Office for Consumer Protection and Food Safety and carried following the guidelines from Directive 2010/63/EU of the European Parliament on the protection of animals used for scientific purposes. The marmosets used in this study included eight animals (*Callithrix jacchus,* five males, and three females). The animals’ age ranged between 3 – 10 years (mean = 5.75 years), and the weight ranged between 382–505 grams (mean = 432.5 grams).

### Anesthesia

Each subject underwent two separate imaging sessions, with a gap of at least two weeks between them. In one session, the marmosets were intubated, mechanically ventilated, and anesthetized with only isoflurane (ISO-only). The med-ISO condition employed a combination of medetomidine infusion and isoflurane inhalation (med-ISO) in self-breathing marmosets.

#### Pilot experiments

The anesthetic concentrations were guided by preliminary pilot studies conducted on four subjects. These initial experiments aimed to identify the minimum isoflurane concentration necessary to sustain sedation during fMRI scanning. We established that a 1.1% concentration was the lowest possible for the ISO-only condition and 0.4% sufficed for the med-ISO condition. We observed that lower concentrations would lead to signs of waking up in the marmosets, characterized by bodily movements, rapidly increasing respiratory rate in self-breathing animals (med-ISO), or breathing no longer synchronized with mechanical ventilation in intubated animals (ISO-only). When these signs were observed, we immediately increased the supplied ISO concentration to sedate the animals.

The medetomidine infusion rate applied in the med-ISO condition was 0.1 mg/kg/h, equivalent to dexmedetomidine at 0.05 mg/kg/h—a dosage typical in rat studies (Sirmpilatze et al., 2019). It’s worth noting that in one of the pilot studies, we experimented with a double rate (0.2 mg/kg/h) of medetomidine infusion. Despite this increase, the subject regained consciousness when the isoflurane concentration dropped below the identified threshold of 0.4%.

During the two sessions included in the study, we opted to use the previously identified threshold ISO concentrations as well as concentrations above the threshold. In the following sections, we describe the anesthetic protocols in detail.

#### Isoflurane-only anesthesia

Isoflurane-only anesthesia (ISO-only condition) was induced with a mixture of alfaxalone (12 mg/kg) and diazepam (3 mg/kg) injected i.m. This was followed by 0.05 ml glycopyrronium bromide per animal (Robinul 0.2 mg/ml, Riemser Biosyn) to prevent secretions, maropitant (1 mg/kg, Cerenia, Pfizer) as an antiemetic, and meloxicam (0.2 mg/kg, Metacam, Boehringer Ingelheim) as an anti-inflammatory analgesic. The animals were then intubated using a custom-made flexible endotracheal tube and mechanically ventilated at 35–37 bpm (Animal Respirator Advanced 4601–2; TSE Systems GmbH, Bad Homburg, Germany). The animals were placed in a prone position inside a custom-built MRI-compatible bed equipped with custom-made stereotaxic apparatus. The ear bars served as hearing protection and were embedded with a lidocaine-containing ointment (EMLA 5%, AstraZeneca) for local anesthesia. A radiofrequency single loop coil with a diameter size of 40 × 43 mm (Rapid Biomedical GmbH, Rimpar, Germany) was fixated on top of the animal’s head. During experiments, dexpanthenol eye ointment (Bepanthen, Bayer) was applied for corneal protection. Monitoring equipment consisted of a rectal temperature probe, a pneumatic pressure sensor attached to the chest, and three surface electrodes for ECG (MR-compatible Model 1030 monitoring and gating system; Small Animal Instruments Inc, Stony Brook, NY 11790, USA). Rectal temperature was kept within 36.5 ± 1°C using a pad filled with circulating heated water. Anesthesia was maintained with isoflurane delivered via the respirator, using a mixture of medical air/O2 (1:1 ratio) as the carrier gas. Each animal’s isoflurane concentration was adjusted to maintain stable anesthesia (range: 1.1 – 1.7%) and physiology. Our goal was to start eah session with 1.4% and end with 1.1%, but in two animals we had to use higher concentrations of up to 1.7%. Each imaging session lasted up to 5 hr, during which multiple anatomical and functional MRI scans were acquired. Resting-state (RS) data were acquired in all 8 animals, consisting of 5 - 9 fMRI runs per session, with each run lasting either 5 or 10 min (370 min of RS fMRI data in total). In 5/8 marmosets we also performed task-based fMRI with visual stimulation. In one animal task-based fMRI was acquired at an isoflurane concentration of 1.4%, in another at 1.1%, and in three animals at both 1.4% and 1.1% concentrations. At each isoflurane concentration level the task-based fMRI runs were repeated 3 times, with each run lasting 5.5 min. At the end of the session, isoflurane was stopped, and the marmosets were extubated as soon as spontaneous breathing was established.

#### Medetomidine anesthesia supplemented with isoflurane

Anesthesia was induced with a bolus intramuscular injection of ketamine 3 mg/kg, medetomidine 0.05 mg/kg, glycopyrrolate 0.01 mg/kg, and maropitant 1 mg/kg. During animal preparation 2% isoflurane was supplied through a mask while an intravenous line (i.v. line) was placed into the vena femoralis for constant infusion. The animal was then placed on a 3D-printed chin plate, and the surface coil was fixed with adhesive tape directly on top of the head. Anesthesia monitoring was performed using the same equipment as for the ISO-only protocol. Before commencing with imaging, the medetomidine i.v. infusion was started at a rate of 0.1 mg/kg/h (in a 1:5 dilution with saline), and the isoflurane concentration was set to 0.6%. An anatomical scan was acquired first, followed by multiple fMRI runs. Specifically the following fMRI runs were performed: 1 resting-state (RS) run lasting 10 min, 3 - 6 task-based fMRI runs with visual stimulation (each lasting 5.5 min) and 1 further RS-fMRI run of 10 min. The medetomidine infusion rate was then decreased to 0.4% and an equilibration period of 30 min was allowed. Subsequently, the same fMRI runs were acquired again at the lower dose (see **Supplementary Figure 13** for a schematic representation). At the end of each imaging session, lasting up to 5 hours, the medetomidine infusion was stopped, isoflurane concentration was lowered to 0% and atipamezole (0.125 - 0.25 mg/kg) was administered intraperitoneally for antagonization.

### Data acquisition

All imaging data were acquired on a 9.4T Bruker BioSpec MRI system, equipped with the B-GA 20S gradient, and operated via ParaVision 6.0.1 software (Bruker BioSpin MRI GmbH, Ettlingen, Germany). Radiofrequency signals were transmitted with a volume resonator (inner diameter 154 mm, Bruker BioSpin MRI GmbH) and received with a 40 × 43 mm loop coil (Rapid Biomedical GmbH, Rimpar, Germany). Before data acquisition, a field map was computed, and shims were adjusted for signal homogeneity in an ellipsoidal volume within the marmoset brain (MAPSHIM). Functional data were collected using a gradient-echo EPI (GRE-EPI) sequence with the following parameters: repetition time 2 sec, echo time 18 ms, flip angle 90°, field of view 62.4 × 25.6 mm2, matrix size 156 × 64 (in-plane resolution of 0.4 × 0.4 mm^2^), 40 contiguous coronal slices with 0.8 mm thickness and no gap. For resting-state runs, we acquired 150 - 300 volumes (5 - 10 min runs), while for visual stimulation, we acquired 165 volumes (5.5 mins runs) with multiple run repetitions per session and anesthetic concentration.

For structural imaging, we acquired a 0.21 mm isotropic magnetization transfer (MT)-weighted volumes using a 3D, RF-spoiled, fast low-angle shot sequence with the following parameters: 2 averages, repetition time 16.1 ms, echo time 3.8 ms, flip angle 5°, the field of view 37.8 × 37.8 × 37.8 mm3, and matrix size 180 × 180 × 180.

### Openly-available awake marmoset resting-state datasets

We obtained awake resting-state datasets from the openly shared resource on the Marmoset Connectome project (https://www.marmosetbrainconnectome.org). We used the NIH data (Sub-06 – Sub-32, n subjects = 26) (Schaeffer et al. 2022), which was acquired using a 7 T 30 cm horizontal bore magnet (Bruker BioSpin Corp, Billerica, MA, USA) with a custom-built 15-cm-diameter gradient coil with 450-mT/m maximum gradient strength (Resonance Research Corp, Billerica, MA, USA) and custom 10-channel phased-array receive coil. Further datasets and acquisition parameters details are available on the Marmoset Connectome Project website (https://www.marmosetbrainconnectome.org).

### Visual stimulation

Visual stimulation was done via a flickering bright LED light source (a single 3W white light LED element) placed at the back end of the magnet bore, about 100 cm away from the monkey’s eyes, and computer-controlled via a TTL pulse. The light flicked at 20 Hz, and the stimulation was presented in five blocks per run (30 baseline, followed by 30 sec ON - 30 sec OFF epochs repeated x5). The eyes of the monkeys were closed and covered with ointment throughout all experiments.

### Data pre-processing

AFNI/SUMA software packages were used for pre-processing all structural and functional imaging data (Cox, 1996). Since data were acquired in the sphinx position, we were required to fix the image plane and orientation before functional analysis. We used a custom-made script, *run/do_afni_reco*, which converted the source data files from DICOM format to NIFTI format using the *Dimon* function in AFNI. The fMRI time series were then corrected for slice-timing differences (‘*3dTshift*’) and despiked (‘*3dDespike*’). To reorient the sphinx position, we used (‘*3dWarp*’) and (‘*3dLRFlip*’) along with (‘*3dresample*’) to fix the original oblique to the cardinal acquisition, shift the Y axis (coronal) and Z axis (axial) orientations and to relabel the header information for the correct orientation (e.g., LPI). Data was then placed in a BIDS format folder organization for further processing.

#### Pre-processing of anatomical data

After fixing the initial sphinx position, we analyzed anatomical data using the *@animal_warper* function in AFNI. The *MB3_v3* marmoset atlas was used as a reference for anatomical segmentation. This script aligns each subject session’s anatomy to the template and stores the warped linear and non-linear transformations. Additionally, the script produces automatic quality control images of the segmentation, which enables the visual inspection of overall alignment results from reference to the source (see **Supplemental Figure 14** for a subject alignment and tissue segmentation example). An affine transformation with 12 degrees of freedom was used for linear alignment, followed by nonlinear warping into the atlas space. Removing non-brain tissue type was performed via skull-stripping of the aligned anatomical data.

#### Pre-processing of functional data

All available data were fully pre-processed via (*‘afni_proc.py’*). Pre-processing blocks for the visual task data included: (*blip align volreg blur mask scale*). The major steps involved EPI distortion correction (e.g., blip-up-blip-down image pairs), motion correction, co-registration of structural and functional data, blurring, and scaling of time series around the mean. EPI image distortion correction (phase-encode-reversed pairs) was performed using two echo times (0° and 180°) entered as input files in *afni_proc* blipping function. The non-linear EPI warp was calculated and applied to each time series (see **Supplemental Figure 15** for example, EPI with and without distortion correction). The volume with minimal root-mean-square deviation over the entire time series was chosen as a reference for motion correction. A rigid body transformation was applied with six motion parameters (3 rotations + 3 translations). Since animals were anesthetized, there was minimal detected motion across sessions and runs. For quality control, *afni_proc* outputs radial correlated estimates to detect potential noise sources (e.g., coil or major movements) that will be reflected on a map showing the local neighborhood correlations. The report also includes a temporal signal-to-noise estimate (TSNR) of the aligned time series (see **Supplemental Figure 16** for an example TSNR estimate of the in-session EPI). The average signal was defined based on the concatenated runs after regression (*all_runs*), while the noise was defined based on the residual signal after regression (*errts*). TSNR was defined as

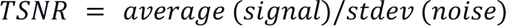

All the EPI volumes were first co-registered to one single reference average EPI volume and then co-registered to the individual in-session anatomical that is aligned to the template reference. Smoothing helped increase the signal-to-noise ratio using a Gaussian kernel of 0.8 mm (double the native resolution of 0.4 mm) to average the signal values among neighboring voxels. For resting-state fMRI data we additionally applied band-pass filtering at 0.005 - 0.12 Hz to remove low-frequency drifts and high-frequency noise. All time series were then scaled by the mean time series, concatenated, and submitted to general linear modeling analyses (GLM) using *3dDeconvolve*. For task-based runs the regression included a gamma block model estimate of the hemodynamic responses function (HRF) lasting 30 seconds for the periods of visual stimulation (5 blocks per run). The resulting estimated beta-coefficients were then used to map areas of significant activation based on a T-statistic scale for significance (see an example of a single subject session activation map in **Supplementary Figure 1**). A cluster activation was defined with a minimum of 50 voxels. For group map results, *3dttest++* was used to perform a student’s t-test on the voxel-wise data for each individual aligned subject 3D beta coefficient map dataset (see **Figure 1**). Functional map visualization was done using AFNI and SUMA for the surface activation mapping. Whole-brain ROI contours based on the *3dMBM_cortex_vM* marmosets atlas were included to highlight active regions on each map.

We considered the necessity of motion regression in our dataset, which entails incorporating six motion parameters and their derivatives as nuisance variables in the GLM. The estimated head motion, quantified as the Euclidean norm of motion derivatives, was typically less than 0.1 mm for most anesthetized scans. To evaluate the impact of motion regression, we conducted all analyses both with and without it and found that its effect was generally minimal. The main manuscript presents results from analyses conducted without motion regression. However, in the supplementary material, we also include some results from analyses with motion regression, particularly when its influence was more pronounced (**Supplementary Figure 5**, **Supplementary Figure 7**, and **Supplementary Figure 10**).

### Visually-evoked BOLD responses in regions-of-interest

Regions-of-interest (ROIs) were defined based on the overlap between the anatomical atlas parcels for LGN, V1, and MT+, and the group-level visual activation map obtained with the med-ISO protocol (see **Supplementary Figure 4 A**).

For visual stimulation fMRI runs acquired with the med-ISO protocol, BOLD signal time series were extracted for each ROI as the mean signal of within-ROI voxels. We then split the extracted time series according to anesthesia concentration — either med-ISO 0.6% or med-ISO 0.4%. Peri-stimulus epochs from −10 sec to +50 sec relative to stimulus onset were extracted, normalized as % signal change relative to the pre-stimulus baseline, and averaged within subjects for each isoflurane level. The analyzed signals resulted in 48 peri-stimulus traces (three ROIs per subject (n = 8) and two anesthetic doses). For each mean peri-stimulus trace, we defined two measures of BOLD response strength — the peak % BOLD signal change and the area under the curve (AUC) during stimulus presentation. To test whether the ISO dose affected the BOLD response strength in the three ROIs, we compared the peak and AUC responses between the two concentrations using paired sample T-tests.

We also applied the same ROIs to visual stimulation fMRI runs acquired during ISO-only anesthesia sessions. Such data were available in 5 out of 8 marmosets (in three subjects at concentrations of both 1.1% and 1.4%, in a fourth subject only at iso 1.1%, and in the fifth one only at iso 1.4%). We repeated the same analysis as for the med-ISO sessions, extracting the mean peri-stimulus traces and computing the peak and AUC BOLD responses for each subject and ROI. This time, however, we skipped the paired samples T-tests between the two isoflurane concentrations since there were only three within-subject paired measurements.

Lastly, we directly compared the two anesthetic protocols, med-ISO and ISO-only. We pooled all visual stimulation fMRI time series for each protocol within the subject (without differentiating between isoflurane concentrations). We visualized the mean peri-stimulus response traces of the three ROIs. We also compared the peak and AUC BOLD responses across the two protocols with paired samples T-tests, relying on the 5/8 marmosets with paired measurements.

### Independent component analysis (ICA)

To reveal resting-state activity patterns, we used independent component analysis (ICA) using the Multivariate Exploratory Linear Optimized Decomposition into Independent Components (MELODIC: http://fsl.fmrib.ox.ac.uk/fsl/fslwiki/MELODIC) of the FSL package. ICA estimates the consistency of spatially and temporally overlapping components over the fMRI time series. Components might consist of meaningful organizing patterns such as the common RSNs and other artifactual effects such as head motion, heart pulsation, or respiration, each carrying an independent spatial pattern and time course. The MELODIC ICA algorithm attempts to segregate the spatial overlap between the components based on the independence of the fMRI-BOLD signals. ICA is a “model-free” algorithm that aims to detect cortical and subcortical responses prevalent among a cluster of voxels instead of the classical modeled BOLD response for comparing the fMRI signal. Previous studies suggested that the optimal number of components lies within the range of 20–30 independent components for RS-fMRI data (Mantini et al., 2012). Here we used 30 components to detect RSNs and submitted the concatenated time series across runs, sessions, and subjects for a grand average ICA map per anesthesia condition - med-ISO, ISO-only, or awake. Condition-based ICA maps were then visualized and thresholded to a significant p-value of < 0.05 and clipped at a maximum Z-score value of (+/−10) for all conditions and components. All ICA analyses were performed twice, with and without implementing motion regression. The impact of motion regression was most noticeable on the awake imaging data (see **supplementary figures 5** and **6**).

### Classification of brain states under isoflurane-only anesthesia

We adapted the PCA-based method from our previous work (Sirmpilatze et al., 2022) to identify burst-suppression during resting-state under ISO-only anesthesia. We modified the analysis in two significant ways. First, we broadened our focus beyond the cortex, extracting the BOLD fMRI signal from a mask that included both cortical and striatal gray matter. This was based on our earlier finding that the striatum is a key component of the burst-suppression network across species, including marmosets (Sirmpilatze et al., 2022). Second, rather than analyzing each fMRI run independently, we performed PCA on the entire concatenated ISO-only resting-state fMRI dataset, which spanned 370 minutes. To ensure continuity and to minimize signal discrepancies at the boundaries of individual fMRI runs, we standardized the voxel time series. This involved subtracting the mean of each run and normalizing by the joint standard deviation across all runs, a normalization technique which is also used in MELODIC ICA when dealing with concatenated runs.

We identified the first temporal principal component (PC1) as indicative of burst-suppression activity, with its peaks representing the hemodynamic correlates of bursts, as previously described (Sirmpilatze et al., 2022). Notably, the peak density in the ISO-only dataset varied significantly, prompting us to use this feature to categorize the data into different states. For peak detection in PC1, we employed a technique commonly used in calcium imaging (Pachitariu et al., 2017): we first smoothed PC1 over time with a Gaussian filter (sigma = 10 seconds) and then applied a rolling maximum of the rolling minimum (window width = 300 seconds). We normalized PC1 by subtracting this baseline and scaling by the maximum signal, placing peaks in a range from 0 (baseline) to 1 (highest peak). Peak identification was done using the *scipy.signal.find_peaks* function in Python’s SciPy library (Virtanen et al., 2020), with parameters set to height 0.15, width 10, distance 4, and rel_height 1.

The entire 370-minute dataset was then divided into 5-minute non-overlapping segments, within which we counted the number of detected peaks. Additionally, we calculated the skewness of the PC1 time series for each segment. We classified these segments into states: ‘no bursts’ (0 peaks), ‘single burst’ (1 peak), and ‘burst-suppression’ (≥ 2 peaks). In one subject (sub-08), burst-suppression was detected in segments with 1.4% isoflurane, but at 1.1%, the clear separation between peaks was lost, which we attributed to the absence of suppression periods. These segments were placed in a separate ‘no suppressions’ category, characterized by ≥ 2 peaks and skew <0.25.

### Network analysis

Networks were constructed by first re-indexing the *MB3_v3* marmoset atlas brain atlas (Liu et al., 2018) to include both the left and the right hemisphere into a single atlas file, including subcortical regions (see **Supplementary Figure 2 B**). The split ROIs included the 106 cortical (*3dMBM_cortex_vH*) as well as 20 subcortical parcels, resulting in a total of 126 parcels per hemisphere (or 252 parcels whole-brain). We obtained the regional time series and connectivity matrices using the program (*3dNetCorr*). Atlas ROIs were down-sampled from the anatomical-based sampling to match the underlying EPI 3D grid (*3dFractionize*), and we then used each ROI to get the mean time series. The time series and adjacency matrices *A*_*ij*_ with n nodes = 252 x 252 were obtained from *3dNetCorr* and were read into Matlab for computing connectivity measures. Subsequently, we mostly relied on the Brain Connectivity Toolbox (BCT) to characterize network properties.

Initially, we calculated network modularity (*Q*) via a hierarchical consensus algorithm. To calculate maximization, we compared the adjacency matrix (*A*_*ij*_) with the expected null connectivity model (*P*_*ij*_) where larger (*Q*) values indicate higher quality. The data-driven approach allows grouping nodes into modules that show high internal density as would maximally be expected from the null model. Consensus modularity (*Q*) is defined as:

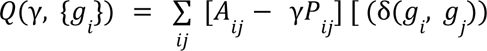

where the γ resolution parameter used for optimization; *g*_*i*∈_{1,…, *C*} is the module assignment of nodes *i* where the Kronecker delta function ઠ _(_*g*_*i*_, *g*_*j*)_ equals one if nodes *i* and *j* belong to the same module *g*_*i*_ = *g_j_*. The function ensures that the total weight of within-module edges is less than the null model. We used the multiresolution consensus function to calculate modularity (Jeub et al., 2018) (https://github.com/LJeub/HierarchicalConsensus).

The community_louvain function from the brain connectivity toolbox was used for the initial community detection (Rubinov and Sporns, 2010) in the functional networks obtained from the openly-available datasets in the awake condition (Schaeffer et al., 2022). All network matrices for all state conditions (e.g., awake, med-ISO 0.6%, med-ISO 0.4%, and ISO-only) were reordered according to the modular organization obtained in the awake condition. The modular structure was obtained on the average hemispheric matrix of the awake condition and then was applied to the whole brain matrix (both hemispheres). The reordering was performed via a desquare/square function, enabling direct comparison across state conditions. To compare the number of modular partitions, we used each individual state and then we plotted the total number of module divisions (**Supplementary Figure 11**). To quantify the differences in ISO-only states we computed the average network strength over all connections and subjects (see **supplementary figure 12**).

Intra- and Inter-hemispheric correlation and distance were evaluated via linear modeling between the left and right hemispheres or across state conditions: awake versus med-ISO or awake versus ISO-only. Additionally, intra- and inter-correlations were evaluated according to their connection distance to observe the distribution of “long-range” and “local” connections and their changes according to the anesthesia state. For additional network measures, we consider that potentially small non-zero values in the matrices may reflect noise measurement rather than an actual correlation (van Wijk et al., 2010). To overcome this potential issue, we thresholded to undirected adjacency matrices based on network density *p*, which is proportional and varies between zero and one, where *p* = 0 indicates no connection available, while *p* = 1 indicates that all connections exist, and 0 < *p* < 1 represents the fraction of all possible connections that are present in the networks. We chose a minimum density value of 0.2 and preserved the overall network structure for each subject and condition.

## Data and Code Accessibility

Imaging data reported in this paper have been shared on the open marmoset connectome project and are available in BIDS format along with all pre-processing scripts. For further information about neuroscience research in non-human primates, please visit the German Primate Center (DPZ) website at (https://www.dpz.eu/de/startseite.html).

## Acknowledgements

We wish to thank Kerstin Fuhrmann and Sina Bode for their help with animal preparation and for the initial experimental preparations.

## Supplementary figures

**supplementary figure 1:**
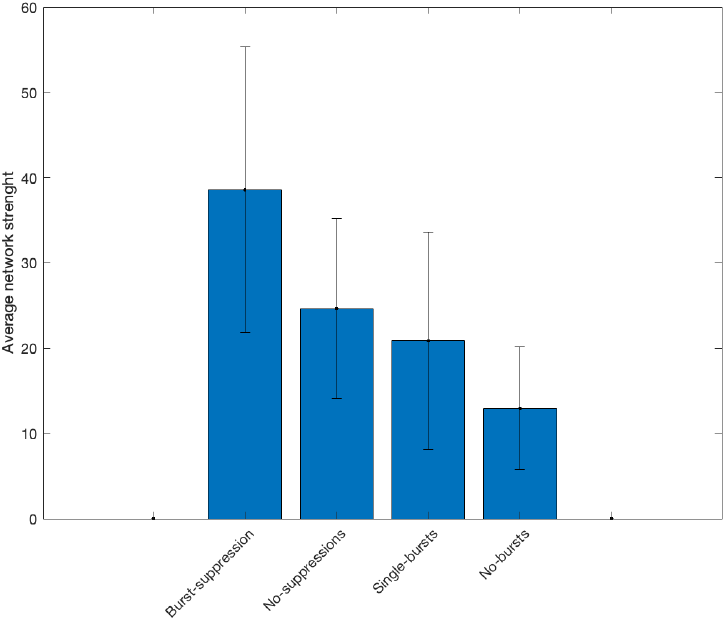
Average strength of network states. For each state under ISO-only anesthesia we calculated the average strength of the network. The strength of the network was four-fold higher for the burst-suppression state as compared to the no-burst state. The decrease in the overall network strength followed the decline in the number of bursts.

**Supplementary Figure 2:**
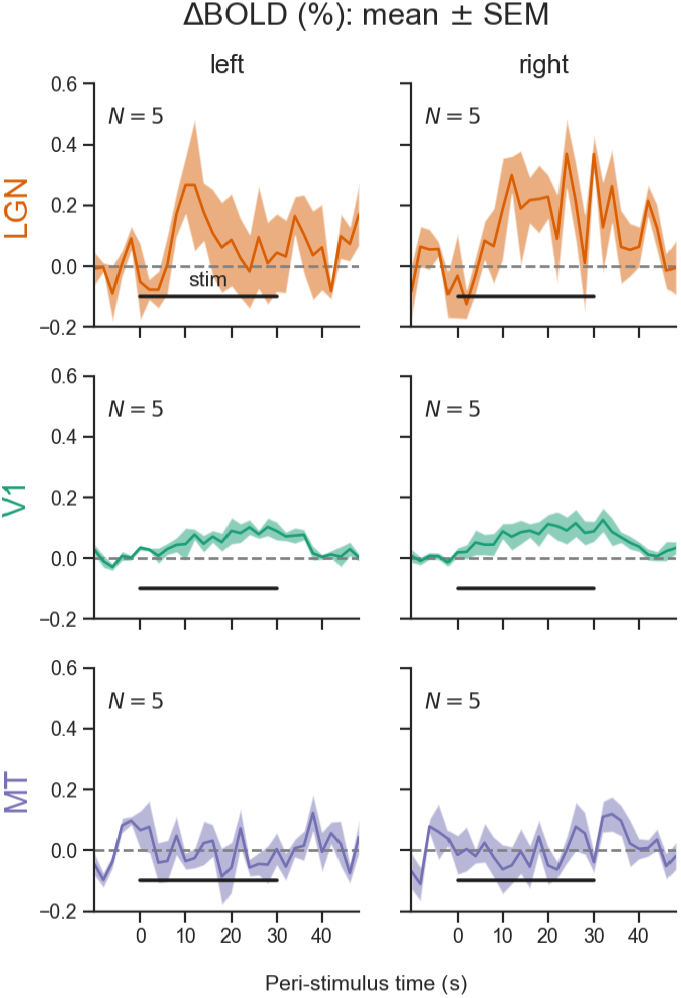
BOLD responses of visual areas during ISO-only anesthesia, shown separately for the left and right hemispheres. Responses to visual stimulation are shown as % BOLD signal change in three regions of interest: LGN, V1, and visual area MT. Visual stimulation blocks were averaged within each subject, separately for the left and right hemispheres. The mean ± SEM response traces are shown across subjects (N = 5). The right LGN showed a more sustained response as compared to the left LGN, which may explain the significant contrast difference between the ISO-only and med-ISO conditions observed in the left hemisphere, but not in the right one (see **Figure 2A** and **Figure 4A**).

**Supplementary Figure 3:**
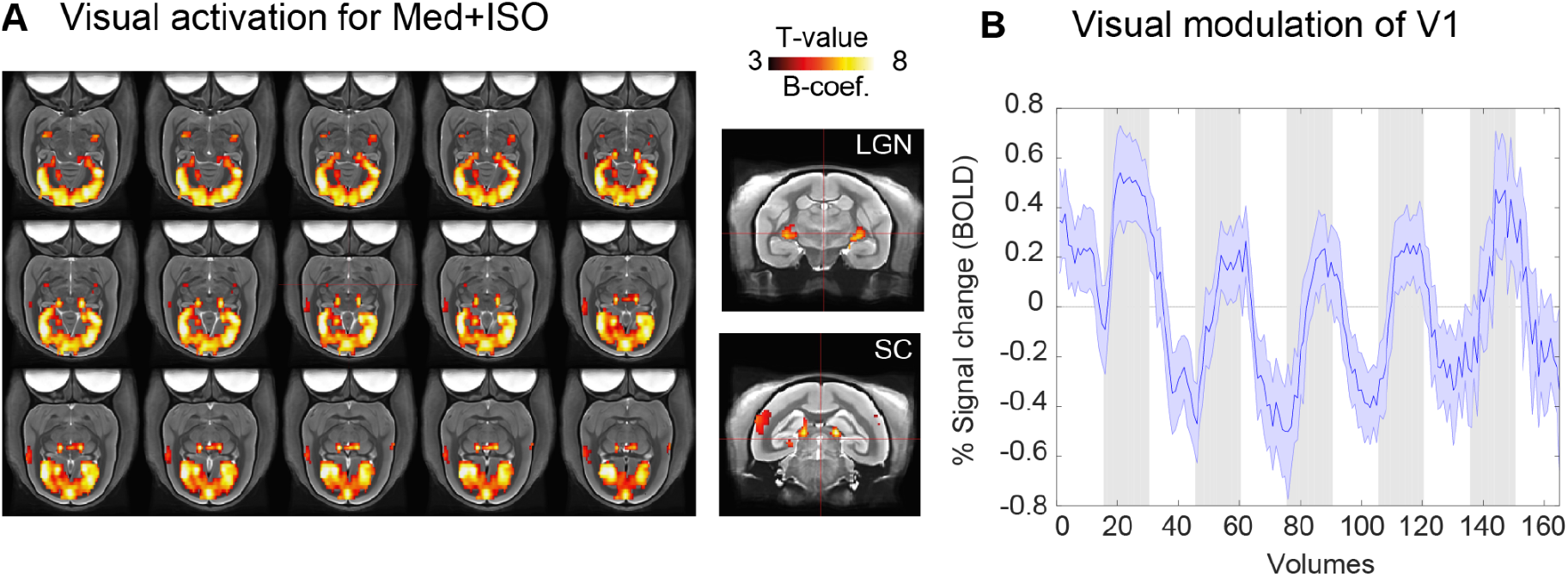
Example activation of a single subject for the med-ISO condition. Activation was found in the LGN, the superior colliculus, the visual cortex, and the motion-sensitive area MT+. **A**.The left panel shows the activation across the regions in the axial plane. The right panels show specific coronal slices that show activation of the LGN and the superior colliculus (SC). **B**. Average signal (mean and ± SEM) time series across active V1 voxels showing five blocks following the visual stimulation rate in gray.

**Supplementary Figure 4:**
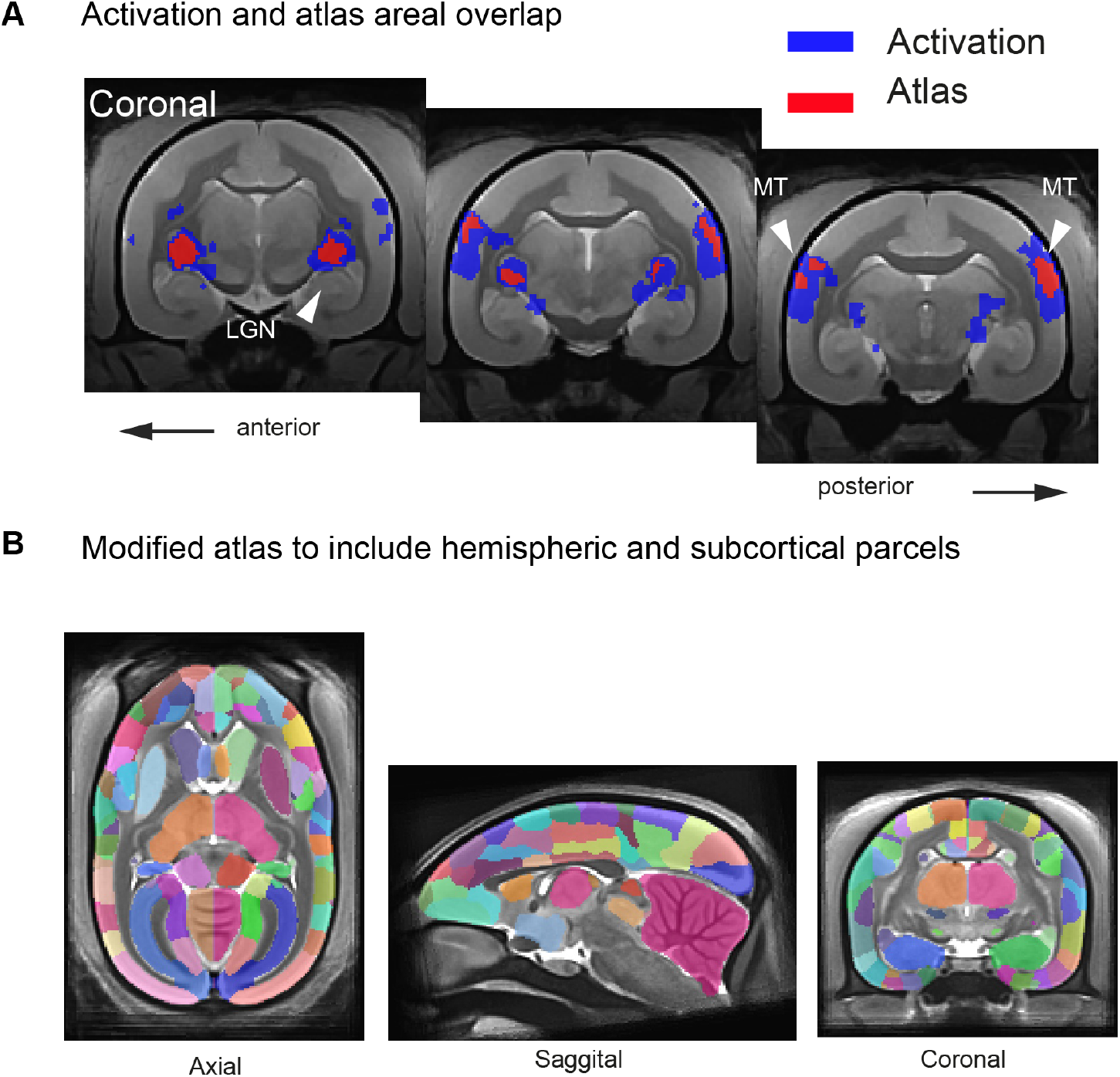
**A.** Overlap areal estimate between visual activation and atlas defined regions-of-interest. Areas with robust activation by visual stimulation during med-ISO anesthesia were found in the lateral geniculate nucleus (LGN), the primary visual cortex (V1), and the area MT. We defined the above three regions of interest using a combination of anatomical and functional activation mapping. Specifically, each ROI was defined as the intersection between the anatomical mask of the respective area (as given by the MBMv3 marmoset atlas) and the functional group-level activation map thresholded at Z-score > 2. Three coronal slices show activation of the LGN and visual motion area MT+. **B.** Atlas parcels were modified to incorporate a continuous count of ROIs from left cortical to left subcortical followed by right cortical and right subcortical. This modification was made for computing matrices that incorporate intra and inter-areal connections.

**Supplementary Figure 5:**
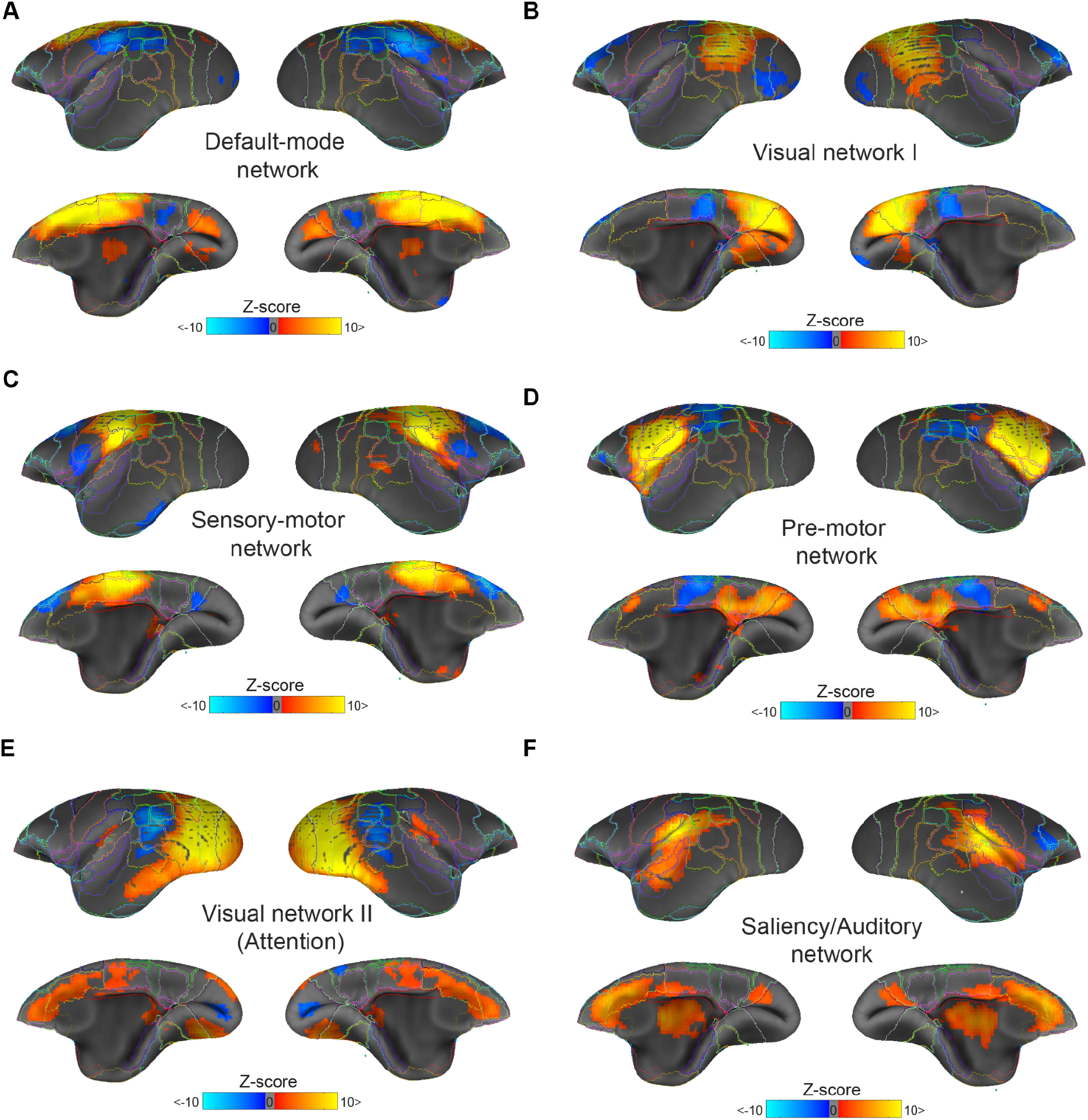
Group-independent component analyses identified common resting-state networks in the awake common marmoset. Analyses were performed without applying motion regression. All networks were thresholded at a p-value < 0.05 and a minimum Z-score of 2 and clipped at a maximum Z-score value of (+/−10). Using these criteria, we identified the following networks: (A) Default-mode network (4.4 % e.v.; 1.69 % t.v.), (B) the sensory-motor network (4.12% e.v.; 1.58 % t.v.), (C) the visual network I (4.27 % e.v.; 1.64 % t.v.), (D) the visual network II (3.81 % e.v.; 1.46 % t.v.), (E) the pre-motor network (3.84 % e.v.; 1.47 % t.v.) and (F) the salience/auditory network (3.69 % e.v.; 1.41 % t.v.).

**Supplementary Figure 6:**
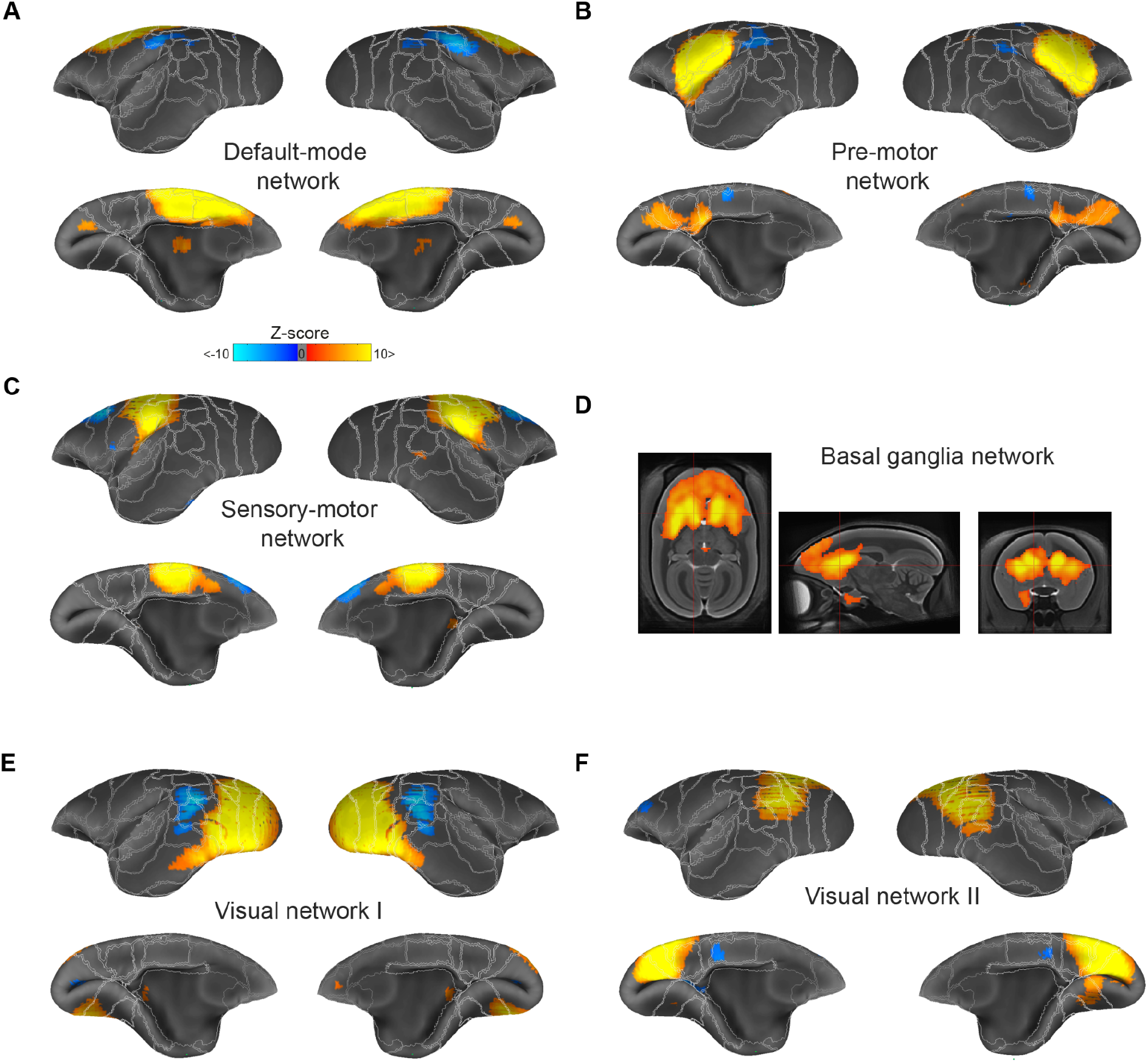
Group-independent component analyses identified common resting-state networks in the awake common marmoset. Analyses were performed with motion regression. All networks were thresholded at a p-value < 0.05 and a minimum Z-score of 2 and clipped at a maximum Z-score value of (+/−10). Using these criteria, we identified the following networks: (A) Default-mode network (4.4 % e.v.; 0.51 % t.v.), (B) the pre-motor network (4.15% e.v.; 0.48 % t.v.), (C) the sensory-motor network (4.31 % e.v.; 0.50 % t.v.), (D) the basal ganglia network (3.33 % e.v.; 0.38 % t.v.), (E) the visual network I (4.23 % e.v.; 0.49 % t.v.) and (F) visual network II (Attention, 4.29 % e.v.; 0.5 % t.v.).

**Supplementary Figure 7:**
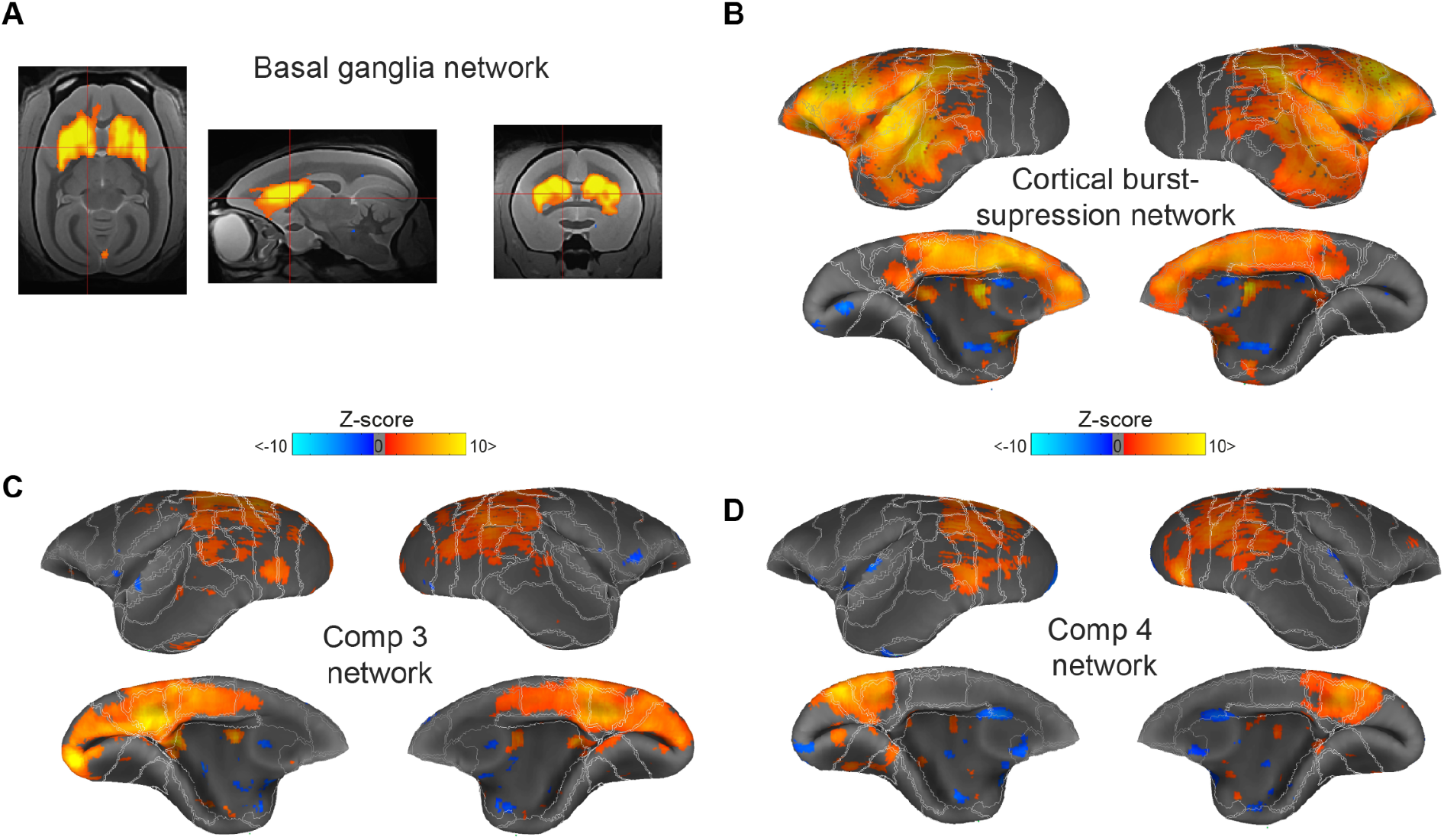
Lack of resting-state network organization under isoflurane-only (ISO-only) anesthesia. Group-independent component analyses identified uncommon networks under ISO-only (1.4 - 1.1 %) anesthesia. All networks were thresholded at a p-value < 0.05 and a minimum Z-score of 2 and clipped at a maximum Z-score value of (+/−10). Using these criteria, we identified the following networks: (A) The basal ganglia network (4.82 % e.v.; 0.32 % t.v.), (B) the cortical burst-suppression network (4.71 % e.v.; 0.32 % t.v.), and (C and D) two additional networks encompassing posterior cingulate and visual cortices (component 3; 3.87 % e.v.; 0.26 % t.v.; component; 3.54 % e.v.; 0.24 % t.v.).

**supplementary figure 8:**
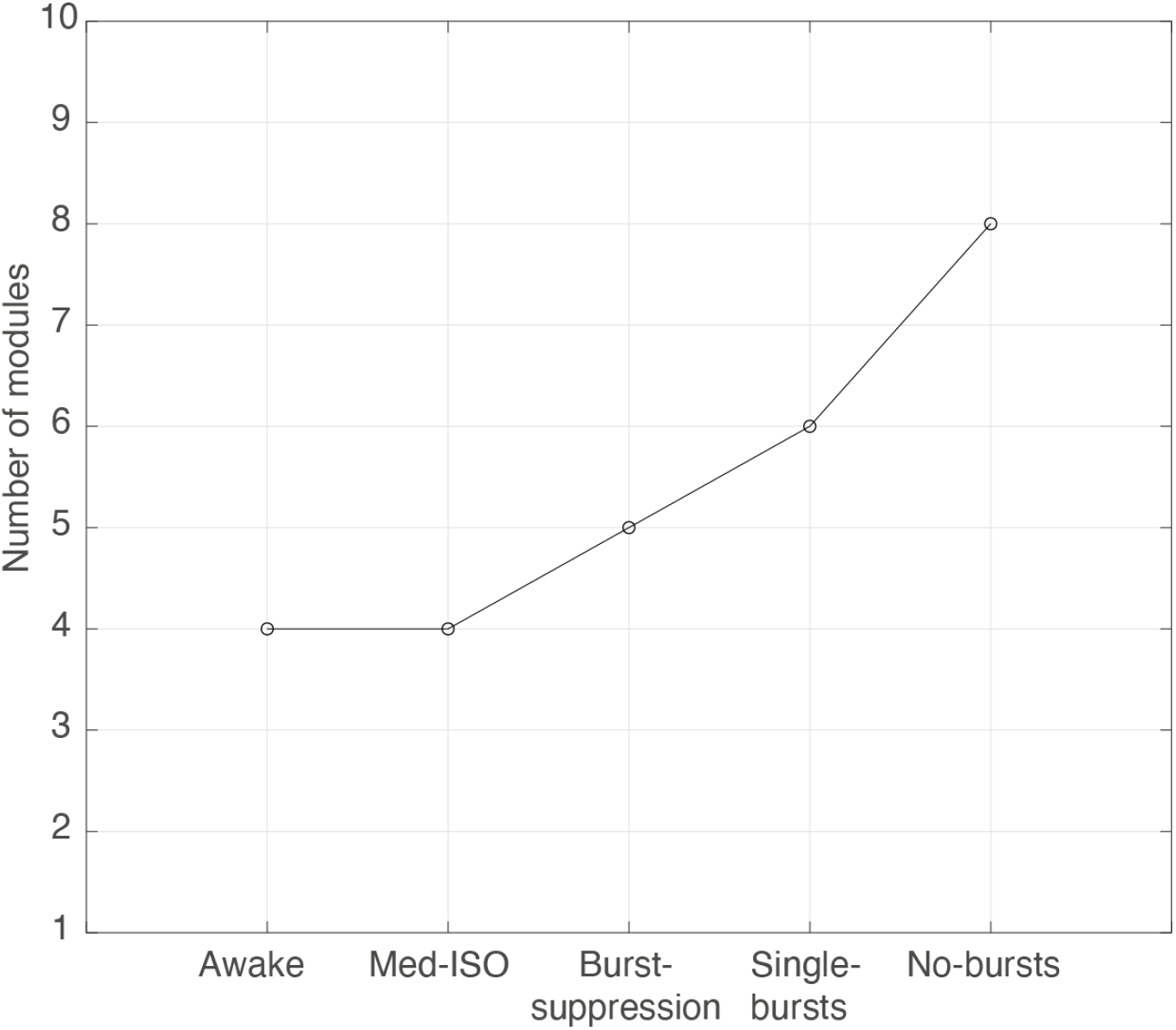
Detected number of modules per state. The Louvain community detection algorithm maximizes the number of within-edges and minimizes the number across groups. The number of modules indicate their maximum of non-overlap groups based on hierarchical modularity obtained for each state: awake, med-ISO, burst-suppression, single-bursts, and no-bursts.

**Supplementary Figure 9:**
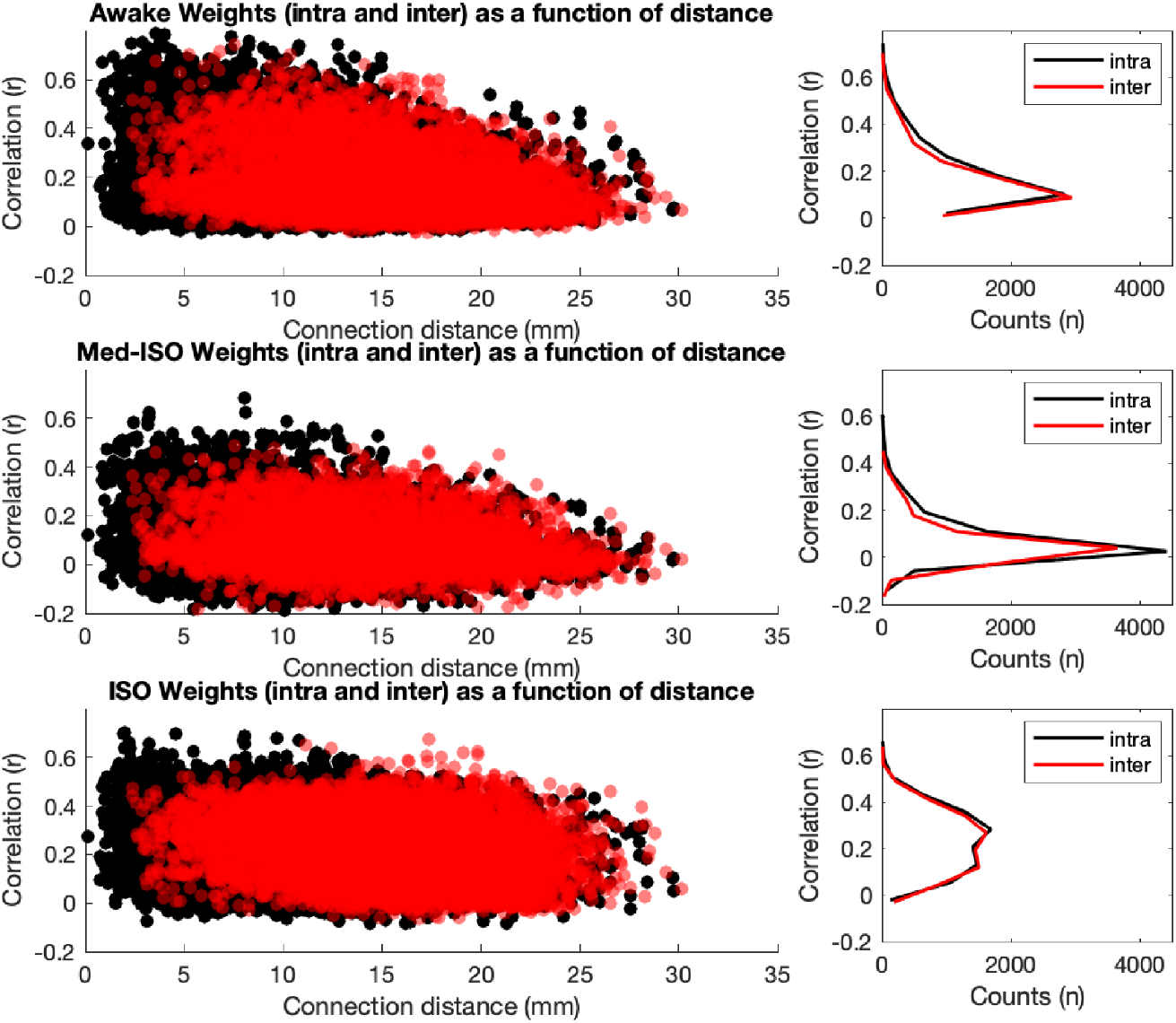
Inter- and intra-hemispheric correlations as a function of distance. Left: Connection weights as a function of node distance for both intra (black) and inter (red) hemispheric correlations and for each state condition, awake (top), med-ISO (center), and ISO-only (bottom). Right: Distribution of intra- and inter-hemispheric correlation coefficients. In contrast to ISO-only, med-ISO revealed a peak shape comparable to the awake state.

**Supplementary Figure 10:**
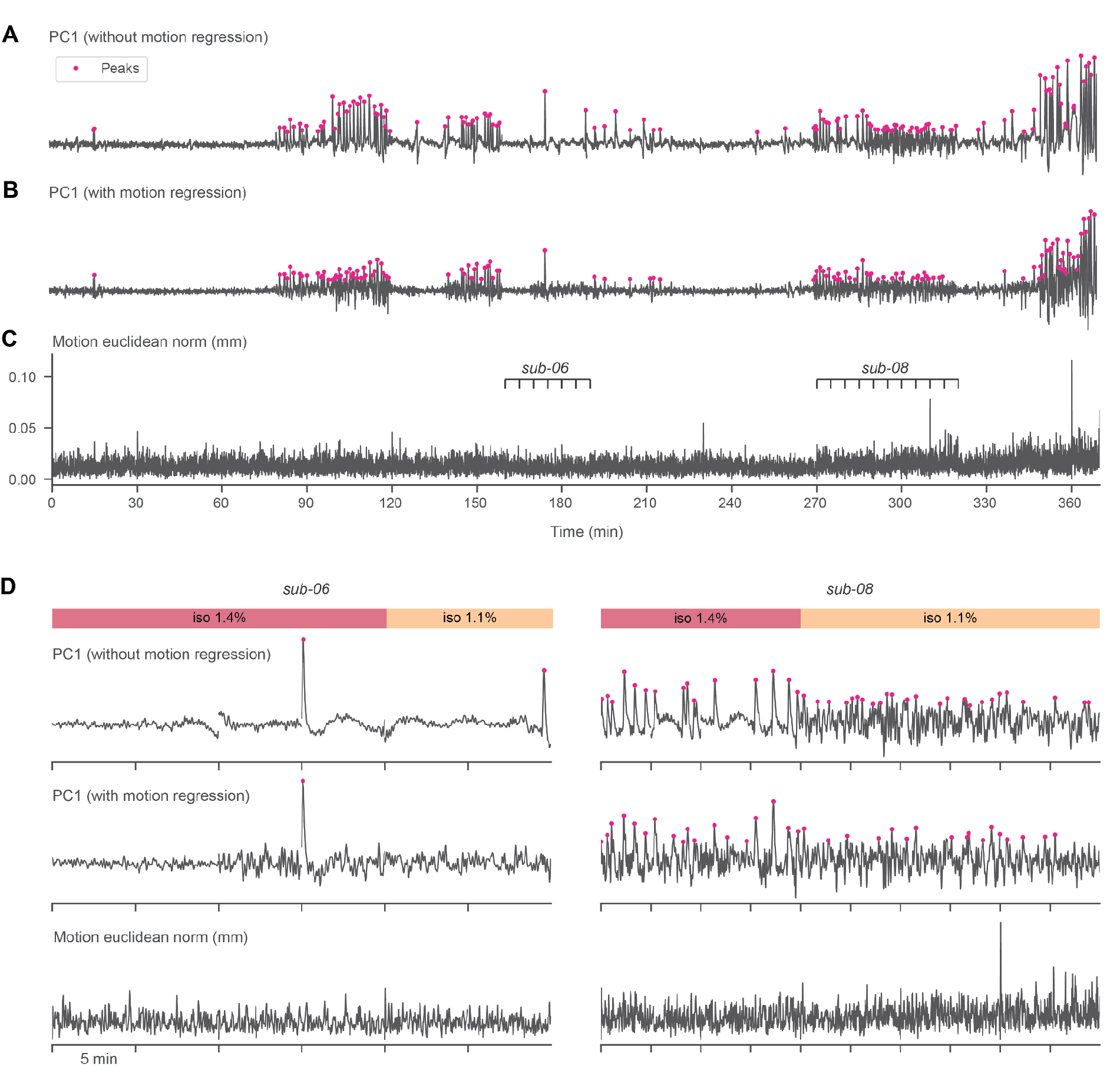
The impact of motion regression on the detection of bursts. **A.** The detrended first temporal Principal Component (PC1) extracted from the cortico-striatal voxels of the concatenated resting-state (RS) fMRI runs acquired with isoflurane-only (ISO-only) anesthesia. The PC1 time series exhibits sharp peaks - presumed to be associated with bursts. This is the same time series as in **Figure 1 C** and is derived from an analysis performed without motion regression. **B.** The same PC1 time series, but derived from an analysis including motion regression. **C.** The estimated head motion during the same period, quantified as the euclidean norm of the motion parameter derivatives. **D**. Examples of two subjects with relatively sparse (sub-06) and dense (sub-08) peaks are highlighted, showing the same three time series: PC1 without motion regression (top), PC1 with motion regression (middle), motion euclidean norm (bottom). This figure highlights that burst peaks are better separated, and therefore easier to detect, when motion is not regressed out.

**supplementary figure 11:**
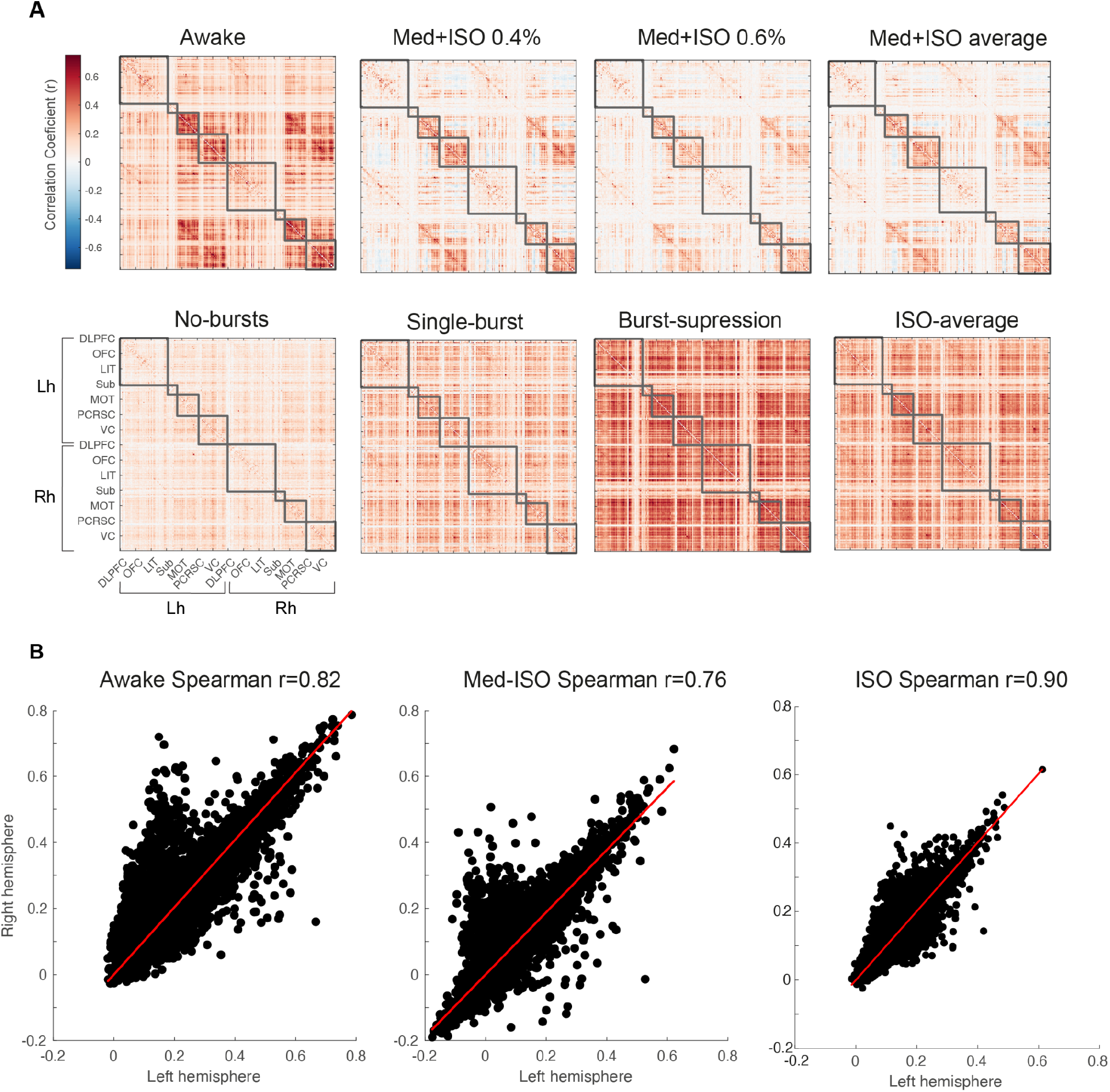
Preservation of functional connectivity structure under med-ISO anesthesia. This figure is analogous to **Figure 6**, but the analyses were performed with motion regression. **A**. Average connectivity matrices for each condition Awake, med-ISO 0.4%, med-ISO 0.6%, and average med-ISO (top). (Bottom) matrices show the ISO-only states for the lack of bursts (no-bursts), the presence of one burst (single-burst), and the clear presence of bursts (burst-suppression), along with the average matrix for all the ISO matrices. The gray squares indicate their maximal no-overlap group based on hierarchical modularity (4 modules per hemisphere) obtained for the awake condition. The Louvain community detection algorithm maximizes the number of within-edges and minimizes the number across groups. The connectivity matrices include each intra-hemispheric correlation and inter-hemispheric correlation. Labels are ordered based on their second-level labeling from the MBV_v3. Labels. The four modules detected largely encompass the frontal cortex (FC), subcortical (Sub), motor cortex (MC), and visual (VC). **B**. Interhemispheric correlation shows the overall connectivity pattern across states; Awake (left), med-ISO (middle), and ISO-only (right). The intersecting line shows the linear fit between the left and right hemispheres. Notice how highly correlated the hemispheres are under the ISO-only condition as compared to the awake and med-ISO conditions.

**Supplementary Figure 12:**
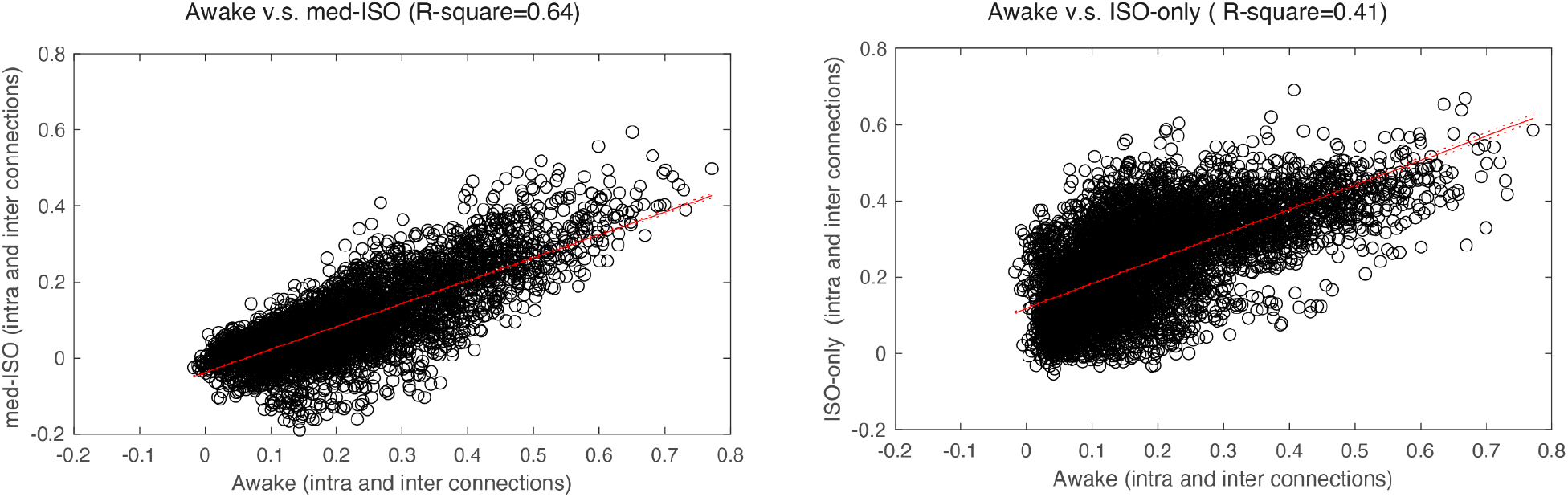
Inter- and intra-hemispheric correlations between awake and anesthetized conditions. **A.** Distribution of connection weights from the awake conditions against the med-ISO condition. The intersecting line shows the linear fit between the awake and med-ISO conditions (Adjusted R-square 0.64). **B.** Distribution of connection weights from the awake conditions against the ISO-only condition (Adjusted R-square 0.41). Notice the more linear trend between the awake v.s. med-ISO condition as compared to the same comparison for the ISO-only condition.

**supplementary figure 13:**
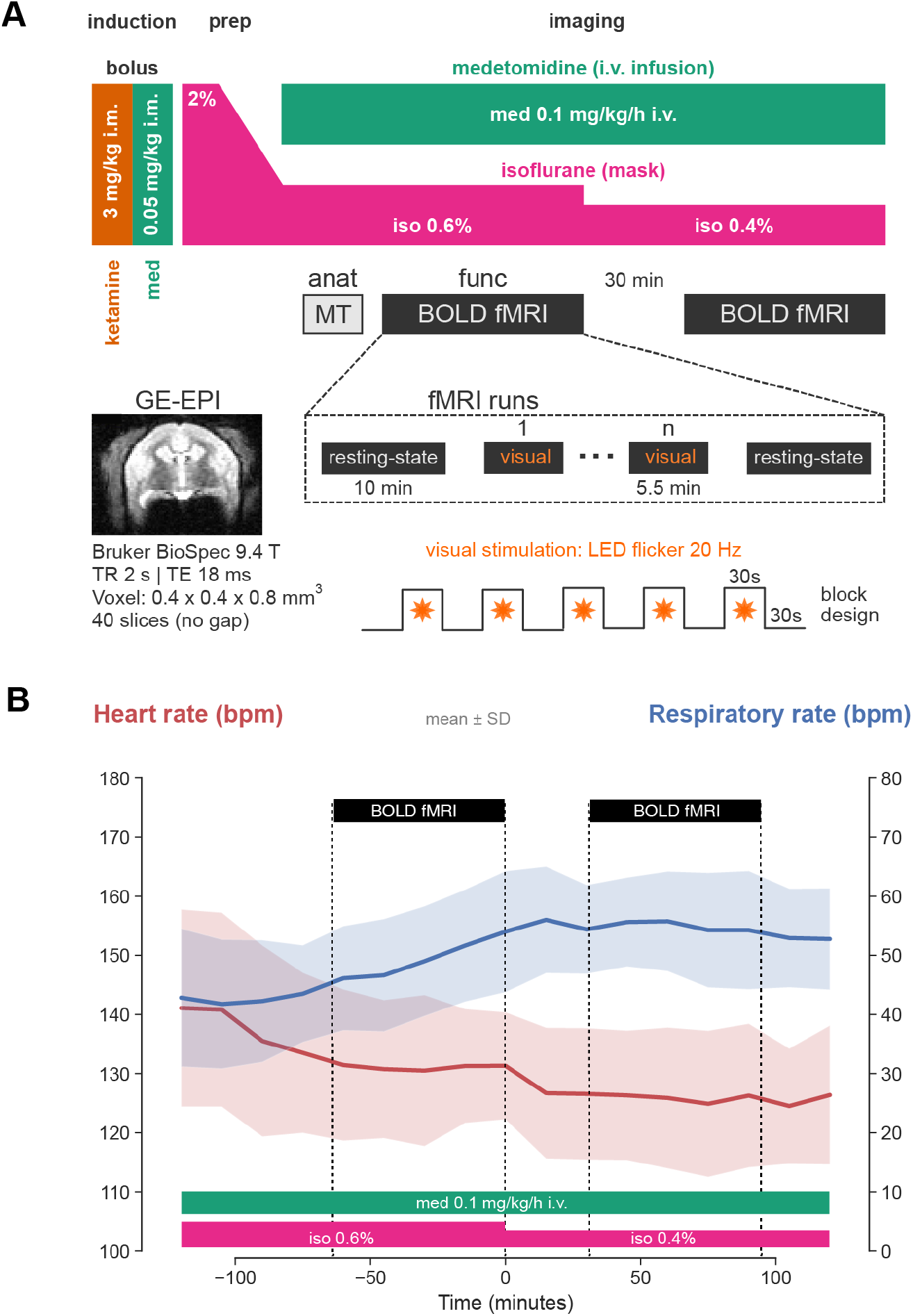
**A.** Anesthetic protocol and data acquisition during med-ISO experiments. Anesthesia was induced with a bolus intramuscular injection of ketamine 3 mg/kg and medetomidine 0.05 mg/kg. Animal preparation was performed under 2% isoflurane delivered through a mask. During imaging, anesthesia was maintained with medetomidine delivered i.v. at a rate of 0.1 mg/kg/h, while isoflurane was reduced to 0.6 %. Data acquisition included an anatomical scan (magnetization transfer MT) acquired at the beginning of the imaging session, followed by functional imaging consisting of resting-state (10 mins) and visual task (5.5 min) runs, with multiple run repetitions (2 resting-state and 3-6 visual task runs). Visual stimulation was performed via a flickering LED light source placed at the end of the magnet bore and delivered at 20 Hz per block (5 blocks, 30 sec each). Echo-planar imaging data were acquired with a 9.4 Tesla system (Bruker BioSpec) with repetition time (TR) of 2 sec, echo time (TE) of 18 ms, and a voxel resolution of 0.4 x 0.4 x 0.8 mm^3^ (40 slices, no gap). The same functional imaging data were also acquired after lowering the isoflurane to 0.4%. **B.** Heart rate and respiration rate (in beats/breaths per minute—bpm) were recorded continuously during the experiment and plotted as mean (solid line) +/− standard deviation (shaded area) across all monkeys. After reducing the isoflurane concentration and starting the continuous medetomidine infusion, the heart rate slightly decreased while the respiration rate increased. Both parameters stabilized over time.

**Supplementary Figure 14:**
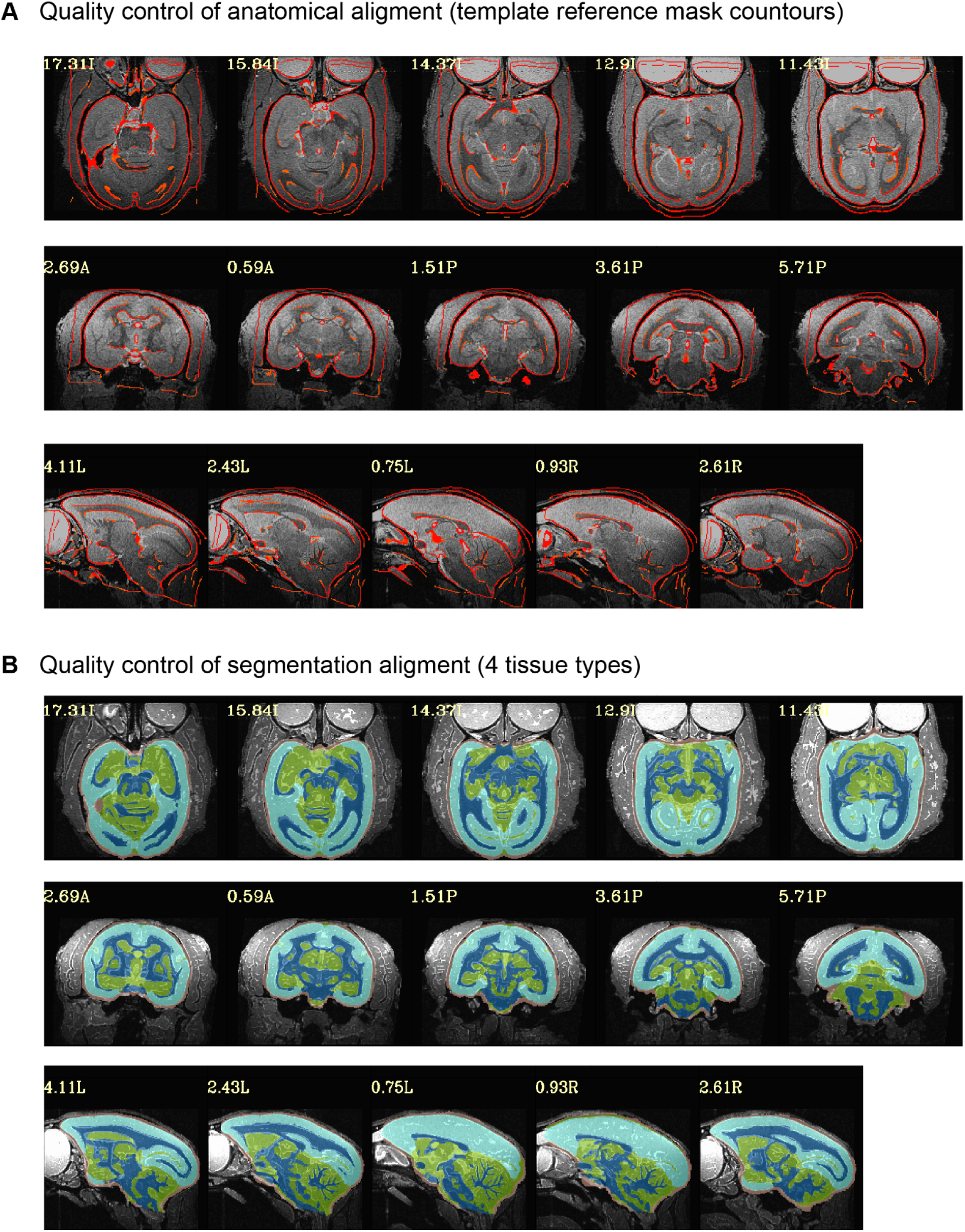
Quality control of linear and non-linear alignment between the in-session anatomy and the reference atlas of an example marmoset subject. **A**. Atlas mask contours (overlay) showing the result of the non-linear alignment of *@animal_warper* AFNI function. Five slices and the respective slice number (upper left) are shown per plane (axial, coronal, and sagittal). The original in-session anatomy is displayed as the underlay. The inverse warp was applied to transform all atlas files into the original scanner space. **B**. Same alignment showing the four segmentation (WM, GM, CSF, and subcortex).

**Supplementary Figure 15:**
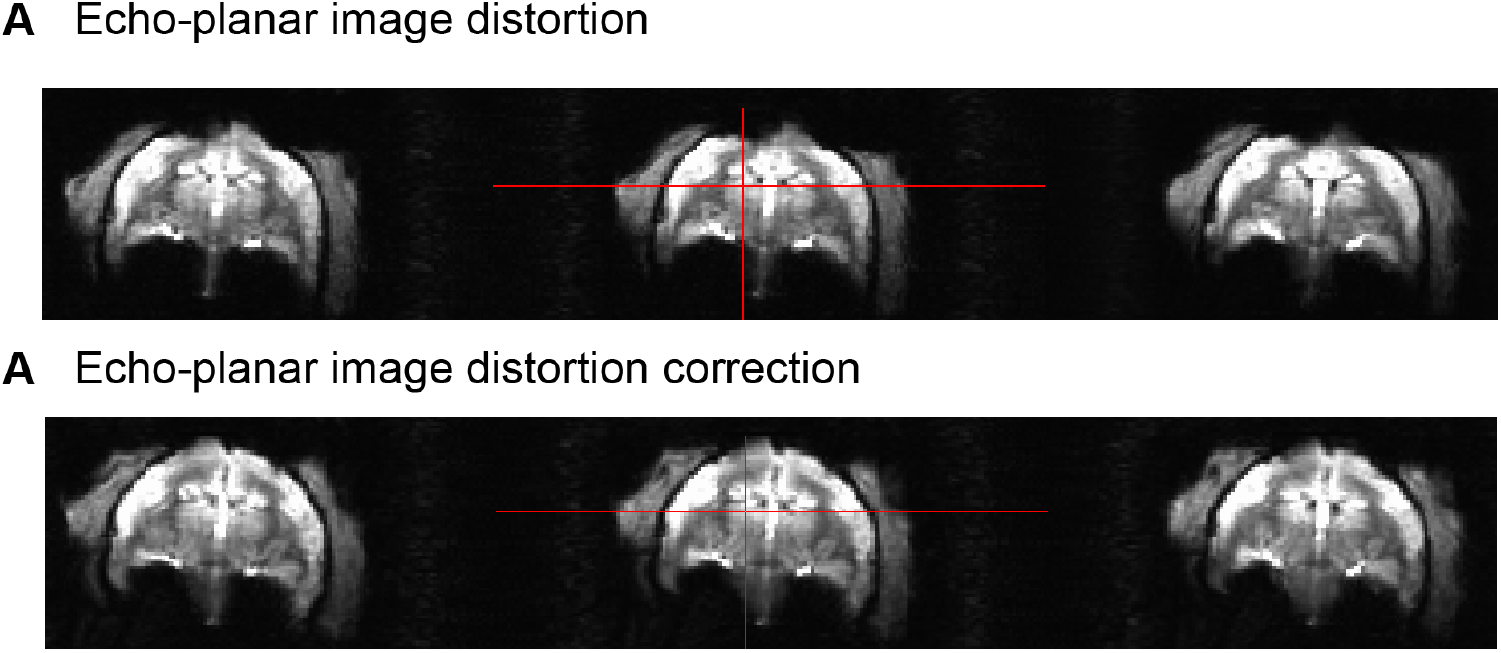
Echo-planar imaging (EPI) distortion correction of an example subject. **A**. Raw EPI images showing the original EPI dataset with distortion at the top of the head near the central sulcus. Cross-hair shows an example center slice where the distortion could be observed at the top of the brain. Prior and subsequent slices also show similar distortion. **B**. Same EPI image slices after distortion correction using two echo times.

**Supplementary Figure 16:**
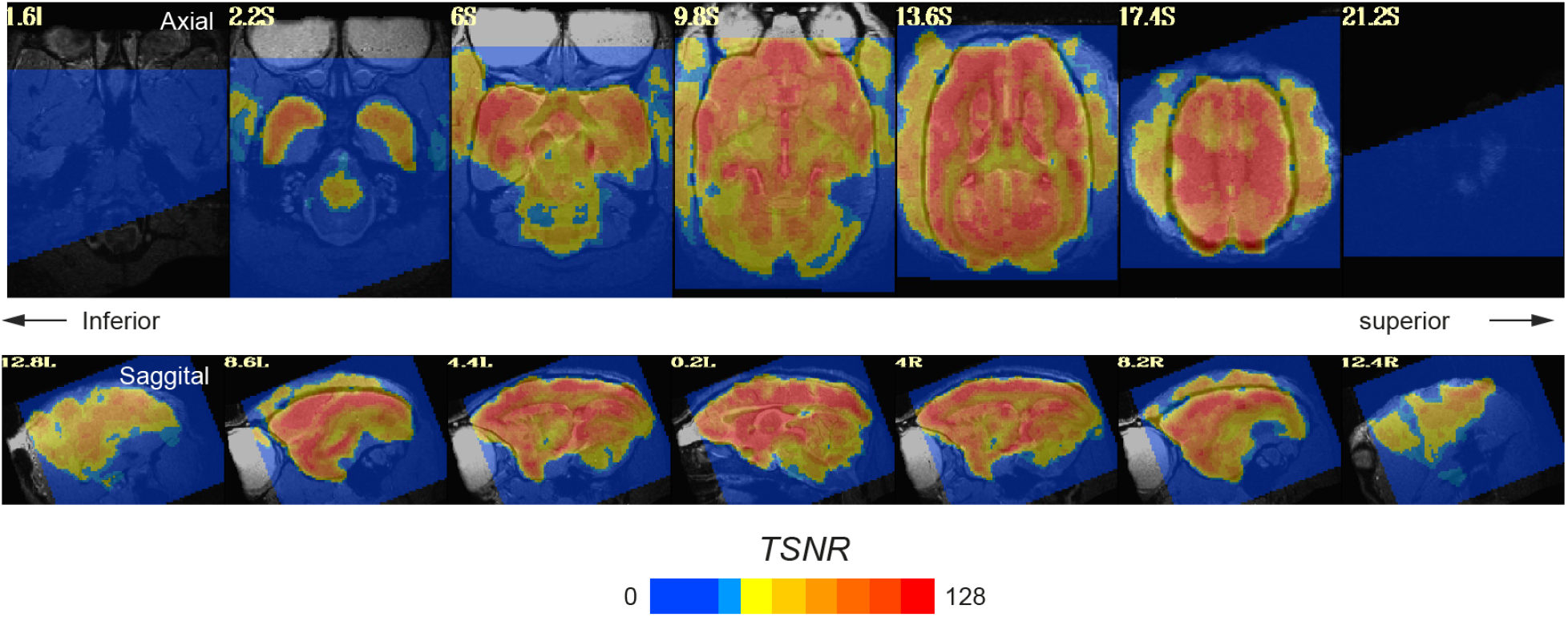
Temporal signal-to-noise ratio (TSNR) of an example subject session. Average TSNR signal overlaid over the anatomical scan with fade image contrast shown for axial and sagittal planes of the aligned dataset. The average signal was defined based on the concatenated runs after regression (*all_runs*). In contrast, the noise was defined based on the residual signal after regression (*errts*). 5 to 95% of the signal lies inside the brain.

